# The Akt-mTOR pathway drives myelin sheath growth by regulating cap-dependent translation

**DOI:** 10.1101/2021.04.12.439555

**Authors:** Karlie N. Fedder-Semmes, Bruce Appel

## Abstract

In the vertebrate central nervous system, oligodendrocytes produce myelin, a specialized proteolipid rich membrane, to insulate and support axons. Individual oligodendrocytes wrap multiple axons with myelin sheaths of variable lengths and thicknesses. Myelin grows at the distal ends of oligodendrocyte processes and multiple lines of work have provided evidence that mRNAs and RNA binding proteins localize to myelin, together supporting a model where local translation controls myelin sheath growth. What signal transduction mechanisms could control this? One strong candidate is the Akt-mTOR pathway, a major cellular signaling hub that coordinates transcription, translation, metabolism, and cytoskeletal organization. Here, using zebrafish as a model system, we found that Akt-mTOR signaling promotes myelin sheath growth and stability during development. Through cell-specific manipulations to oligodendrocytes, we show that the Akt-mTOR pathway drives cap-dependent translation to promote myelination and that restoration of cap-dependent translation is sufficient to rescue myelin deficits in mTOR loss-of-function animals. Moreover, an mTOR-dependent translational regulator co-localized with mRNA encoding a canonically myelin-translated protein in vivo and bioinformatic investigation revealed numerous putative translational targets in the myelin transcriptome. Together, these data raise the possibility that Akt-mTOR signaling in nascent myelin sheaths promotes sheath growth via translation of myelin-resident mRNAs during development.

**SIGNIFICANCE STATEMENT:** In the brain and spinal cord oligodendrocytes extend processes that tightly wrap axons with myelin, a protein and lipid rich membrane that increases electrical impulses and provides trophic support. Myelin membrane grows dramatically following initial axon wrapping in a process that demands protein and lipid synthesis. How protein and lipid synthesis is coordinated with the need for myelin to be generated in certain locations remains unknown. Our study reveals that the Akt-mTOR signaling pathway promotes myelin sheath growth by regulating protein translation. Because we found translational regulators of the Akt-mTOR pathway in myelin, our data raise the possibility Akt-mTOR activity regulates translation in myelin sheaths to deliver myelin on demand to the places it is needed.

## INTRODUCTION

During development, oligodendrocytes extend numerous processes that survey the environment and ensheath target axons with myelin. Notably, not all myelin sheaths produced by an individual oligodendrocyte are uniform in length and thickness (Murtie et al., 2007; Almeida et al., 2011; Chong et al., 2012) and changes in neural activity can influence the amount of myelin made on an axon (Hines et al., 2015; Koudelka et al., 2016; Mitew et al., 2018). Together, this suggests a mechanism whereby oligodendrocytes use localized cell signaling events to coordinate myelin production at distal locations with extracellular cues.

One strong candidate to mediate distal myelin production is the PI3K-Akt-mTOR signaling pathway, whose precise regulation is critical for proper myelin formation. When pathway activity was increased through either oligodendrocyte specific deletion of the antagonist phosphatase PTEN (Goebbels et al., 2010; Harrington et al., 2010) or overexpression of constitutively active Akt (caAkt) (Flores et al., 2008) oligodendrocytes made thicker myelin. Chronic treatment of mice overexpressing caAkt in oligodendrocytes with rapamycin, an mTOR inhibitor, reduced hypermyelination to wild type levels (Narayanan et al., 2009), and decreased pathway activity resulting from oligodendrocyte-specific knockout of mTOR or Raptor, the defining protein of the mTORC1 complex, resulted in severe hypomyelination of the spinal cord (Bercury et al., 2014; Lebrun-Julien et al., 2014; Wahl et al., 2014). Additionally, animals harboring a mutation in the ubiquitin ligase Fbxw7, which targets mTOR for degradation (Mao et al., 2008), made longer myelin sheaths in an mTOR-dependent manner (Kearns et al., 2015).

Protein translation and myelin protein synthesis are among the numerous cell biological processes regulated by the Akt-mTOR pathway. One way that Akt-mTOR signaling regulates protein translation is through the eukaryotic translation initiation factor 4E binding protein (4E-BP) family of proteins. Unphosphorylated 4E-BPs inhibit cap-dependent translation initiation by binding to eIF4E on the 5’ cap of mRNAs (Pause et al., 1994). Upon phosphorylation by the mTORC1 subcomplex, 4E-BP1 dissociates from its binding partner eIF4E, thereby allowing for recruitment of eIF4A and eIF4G to assemble the eIF4F translation initiation complex and promotion of translation (Pause et al., 1994). Dysregulation of Akt-mTOR signaling in oligodendrocytes results in aberrant levels of some myelin proteins (Flores et al., 2008; Narayanan et al., 2009; Bercury et al., 2014; Lebrun-Julien et al., 2014; Wahl et al., 2014; Zou et al., 2014). In vitro, myelin basic protein mRNA is packaged into granules and actively transported into some but not all myelin sheaths (Ainger et al., 1993; Ainger et al., 1997), where it is presumably translated. In support of this possibility, free ribosomes have been observed by electron microscopy in the distal ends of cultured oligodendrocytes (Lunn et al., 1997) and recently hundreds of other myelin resident mRNAs have been identified (Thakurela et al., 2016) and a subset validated in vivo (Yergert et al., 2021).

Here, using zebrafish as a model system, we directly tested the hypothesis that protein translation downstream of Akt-mTOR signaling promotes myelin development. We found that modulation of the Akt-mTOR signaling pathway controlled myelin sheath length and stability. Oligodendrocytes co-localized mRNA and translational regulators downstream of mTOR in myelin, and we identified hundreds of putative mTOR translational targets in myelin. Finally, we performed cell-specific manipulation of protein translation and found that Akt-mTOR signaling required cap-dependent translation to promote proper myelin development. Together, our data raise a model wherein mTOR signaling in myelin sheaths regulate sheath growth by promoting translation of myelin-resident mRNAs.

## METHODS

### CONTACT FOR REAGENT AND RESOURCE SHARING

Further information and requests for resources and reagents should be directed to and will be fulfilled by the Lead Contact, Bruce Appel.

### EXPERIMENTAL MODEL AND SUBJECT DETAILS

#### Zebrafish lines and husbandry

All procedures were approved by the University of Colorado Anschutz Medical Campus Institutional Animal Care and Use Committee (IACUC) and performed to their standards. Wild-type non-transgenic embryos were generated by crosses of male and females of the AB strain. *Tg(sox10:mRFP)* embryos were generated by outcrossing *Tg(sox10:mRFP)^vu234^* (Kucenas et al., 2008) carriers with wild-type AB fish. *mtor* mutant larvae were generated by incrosses of heterozygote *mtor^xu015Gt^* (Ding et al., 2011) carriers. Embryos and larvae were raised at 28.5°C in E3 media (5mM NaCl, 0.17mM KCl, 0.33 mM CaCl_2_, 0.33 mM MgSO_4_ at pH 7.4 with sodium bicarbonate) and staged according to hours and days postfertilization. Only larvae displaying good health and normal developmental patterns were used experimentally. Animals used were of indeterminate sex, as sexual differentiation is not apparent until later larval ages.

### METHOD DETAILS

#### Genotyping

*mtor* mutant larvae were genotyped following imaging. Whole anesthetized larvae were lysed by incubation in 50mM NaOH at 95°C for 30 minutes followed by neutralization with 1/10^th^ volume Tris pH 8.0. PCR amplification was performed using a small volume of lysis and the primers: forward primer: 5′-ATAAGAAAAGAAACCACATGTCATACC-3′; reverse primer: 5′-CTTACCACTCAGAGAGACCAAAG-3′; 5′LTR primer: 5′-CCCTAAGTACTTGTACTTTCACTTG-3′.

#### Plasmids and Expression Construct Generation

Expression constructs were generated by Multisite Gateway cloning using LR Clonase II Plus (Thermo Fisher Scientific) to recombine entry clones p5E, pME, and p3E into a destination vector, pDEST. All entry clones and expression constructs were screened by restriction digestion and sequence confirmed by Sanger sequencing. The following entry clones were used in this study:

p5E: *p5E-sox10; p5E-mbpa*

pME: *pME-mNeonCAAX; pME-myrAkt; pME-4EBP1; pME-4EBP1^T37/46A^; pME-mScarletCAAX; pME-eIF4E; pME-mbpaCDS-24xMBS; pME-HA-NLS-tdMCP-tagRFP* p3E: *p3E-P2A-mNeonCAAX; p3E-eGFP; p3E-P2A-myrAkt; p3E-polyA; p3E-mbpa-3’UTR*

pDEST: *pDEST-Tol2-pA*

*pcDNA3-TORCAR* (Addgene Plasmid 64927) (Zhou et al., 2015) and *pcDNA3-TORCAR(T/A)* (Addgene Plasmid 64928) (Zhou et al., 2015) were gifts from Jin Zhang and used to create *pME-4EBP1* and *pME-4EBP1^T37/46A^*, respectively. *pHA-eIF4E* (Addgene Plasmid 17343) (Okumura et al., 2007) used to generate pME-eIF4E was a gift from Dong-Er Zhang. *pme-HA-NLS-tdMCP-tagRFP* (Addgene Plasmid 86244) (Campbell et al., 2015) was a gift from Florence Marlow.

#### Table of DNA oligonucleotides used for plasmid creation

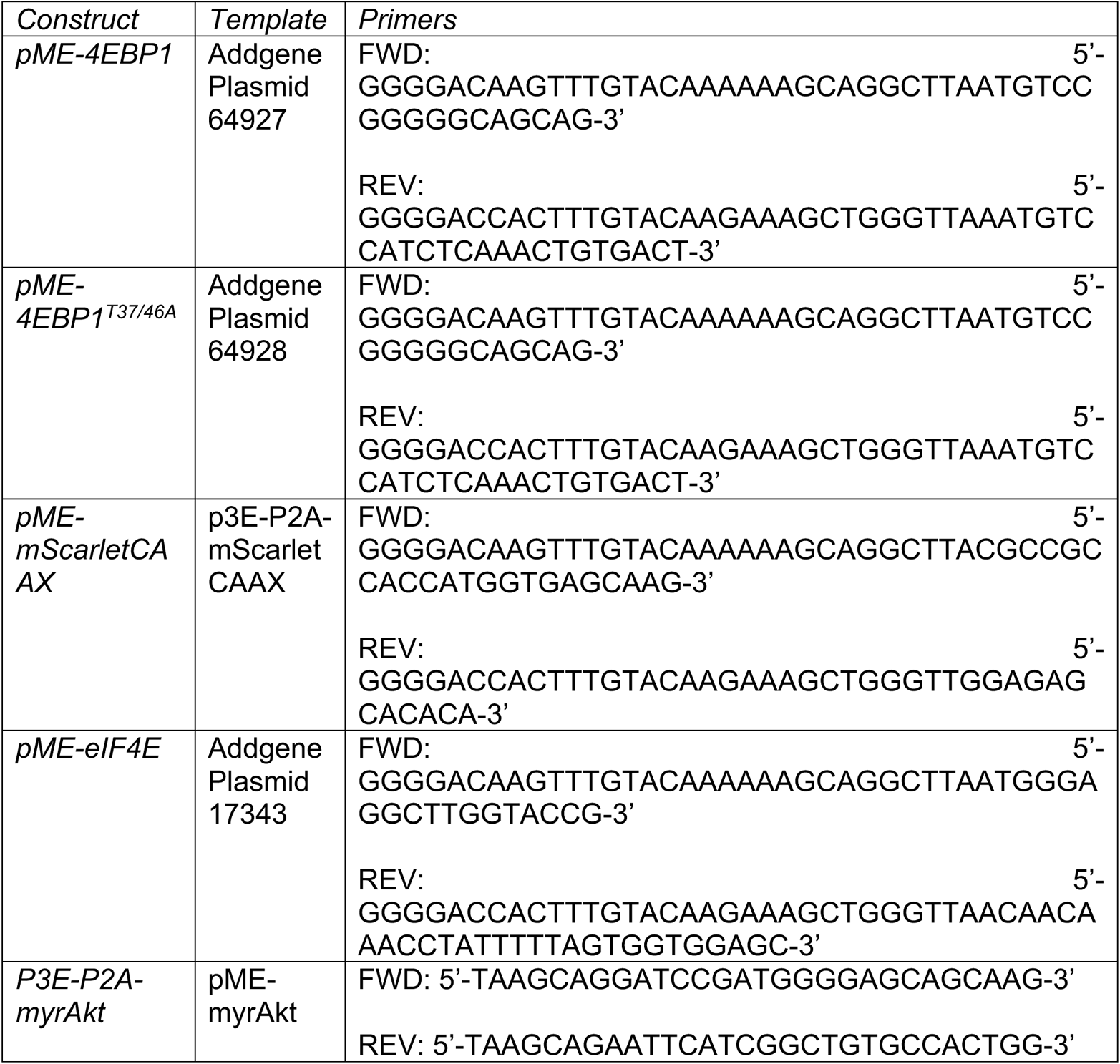

#### Microinjections

Microinjections were performed into newly fertilized eggs generated by incrosses of wild-type AB or heterozygote *mtor^xu015Gt^* carriers. Injection solutions contained 5 µL 0.4M KCl, 250ng Tol2 mRNA, a total of 250ng plasmid DNA, and water to 10 µL. In instances where multiple plasmids were injected, equal concentrations of each plasmid were used. Larvae were raised to the developmental timepoint indicated in individual experiments and selected for good health before inclusion in experiments.

#### Microscopy

For static imaging experiments, larvae were anesthetized with 0.06% tricaine and immobilized in 0.6% low-melt agarose with 0.4% tricaine. For dynamic live imaging experiments, larvae were immobilized in 0.3 mg/ml pancuronium bromide aided by a small tail slit and embedded in 1.2% low-melt agarose. For analysis of myelin sheath length and dynamics, we acquired images using a C-Apochromat 63x, 1.20 NA water immersion objective on a Zeiss CellObserver Spinning Disk confocal microscope equipped with a Photometrics Prime 95B camera. For analysis of subcellular localization, we acquired images using a Plan-Apochromat 63x, 1.4 NA oil immersion objective on a Zeiss LSM 880. Images were collected with Zen (Carl Zeiss), blinded, and processed/analyzed in ImageJ/Fiji (Schindelin et al., 2012).

### QUANTIFICATION AND STATISTICAL ANALYSIS

#### Quantification of myelin sheath length and number

Three dimensional images of single spinal cord oligodendrocytes were collected in Zen. We then used the Fiji plugin Simple Neurite Tracer (Longair et al., 2011) to quantify myelin sheath length and number. Briefly, blinded 8-bit z-stack images of single oligodendrocytes were opened in Simple Neurite Tracer. We traced each sheath and saved the path length (sheath length), path number (sheath number), and the sum of the path lengths (cumulative sheath length) in Excel for further analysis in R and Prism 9.

#### Quantification of myelin sheath dynamics

Three dimensional images of the spinal cord were collected every 15 minutes for 4 hours in Zen. Z-volume movies collected from individual larvae were opened in Fiji. We used the trace segmented line tool to trace individual myelin sheaths. Single myelin sheaths were measured in all frames and exported to Excel before continuing to the next. Myelin sheath dynamic data was then transferred to Prism 9 for visualization and analysis.

#### Colocalization quantification

Three dimensional images of spinal cord oligodendrocytes co-expressing *sox10:4E-BP1-eGFP* and *mbpa* MS2 reporter constructs were collected in Zen. We imported these images to Fiji and used JACoP (Just Another Colocalization Plugin) (Bolte and Cordelières, 2006) to quantify colocalization. Specifically, we isolated single optical planes of individual myelin sheaths and used JACoP to calculate Mander’s Colocalization Coefficients, M1 and M2. M1 and M2 describe the proportion of protein 1 that colocalizes with protein 2 and vice versa, independent of fluorescent intensity. These data were saved in Excel and exported to Prism 9 for visualization and analysis.

#### Statistics

Plots were generated in R (version 3.6.3) with RStudio using ggplot2 (Wickham, 2016) or biomaRt (Durinck et al., 2009) or Prism 9. All statistical tests were performed in Prism 9. We first tested normality of data. In no instances were all datasets normal. For unpaired comparisons, we used Mann-Whitney test to compare ranks. For multiple comparisons, we first assessed overall significance with the Kruskal-Wallis test. If the Kruskal-Wallis test showed significance, we then performed pairwise Mann-Whitney tests with Bonferroni-Holm correction for multiple comparisons.

### BIOINFORMATIC ANALYSIS

#### FIMO analysis

We used Find Individual Motif Occurrences (FIMO) (version 5.3.3), part of the MEME suite software, to identify transcripts containing TOP-like or CERT motifs in the myelin transcriptome. We downloaded the 5’UTR sequences from the myelin transcriptome and created position weight matrices corresponding to TOP-like or CERT motifs. We uploaded the 5’UTR sequences and position weight matrices to FIMO. We changed options to scan the given strand only and otherwise used default settings. We downloaded the resulting file and removed duplicates to generate the final list of unique genes containing TOP-like or CERT motifs.

#### Gene ontology analysis

We used DAVID software (version 6.8) to identify gene ontology terms associated with transcripts containing TOP-like or CERT motifs. We submitted Ensembl Gene IDs of transcripts identified as having these motifs and selected categories for biological process, cellular compartment, and up_keywords. We filtered for false discovery rate less than 0.05 and removed any duplicate terms. We sorted terms from lowest to highest false discovery rate and selected the top 20 for analysis.

## RESULTS

### The Akt-mTOR pathway drives myelin sheath growth

We previously showed that homozygous *mtor* mutant zebrafish larvae express abnormally low levels of myelin genes (Kearns et al., 2015), consistent with studies from mice showing that conditional loss of mTOR function in oligodendrocytes results in a decreased expression of some myelin genes and proteins and decreased myelin sheath thickness (Wahl et al., 2014). For this study, we assessed the number and length of myelin sheaths formed by oligodendrocytes as a sensitive measure of how mTOR activity promotes myelin development. To label individual oligodendrocytes, we used Tol2-mediated transgenesis (Kwan et al., 2007) and *sox10* regulatory DNA to express the membrane localized fluorophore mNeonCAAX in oligodendrocyte lineage cells. We injected *sox10:mNeonCAAX* into newly fertilized eggs generated from incrosses of wild-type or *mtor* heterozygous zebrafish to generate homozygous *mtor* mutant loss-of-function animals. At 5 days post-fertilization (dpf) we imaged spinal cord oligodendrocytes (Figure 1B) and analyzed oligodendrocyte morphology. On average, myelin sheaths in *mtor* mutant larvae were ~35% shorter than those in wild-type larvae (Figure 1C). Additionally, oligodendrocytes in *mtor* mutant larvae produced fewer sheaths on average than those in wild-type larvae (Figure 1D). Together, this reduction of both myelin sheath length and number resulted in a nearly 50% reduction in the cumulative length of myelin produced by individual oligodendrocytes of *mtor* mutants when compared to wild-type controls (Figure 1E). These data therefore show that mTOR function promotes the length and number of myelin sheaths formed by spinal cord oligodendrocytes.

**Figure 1.**
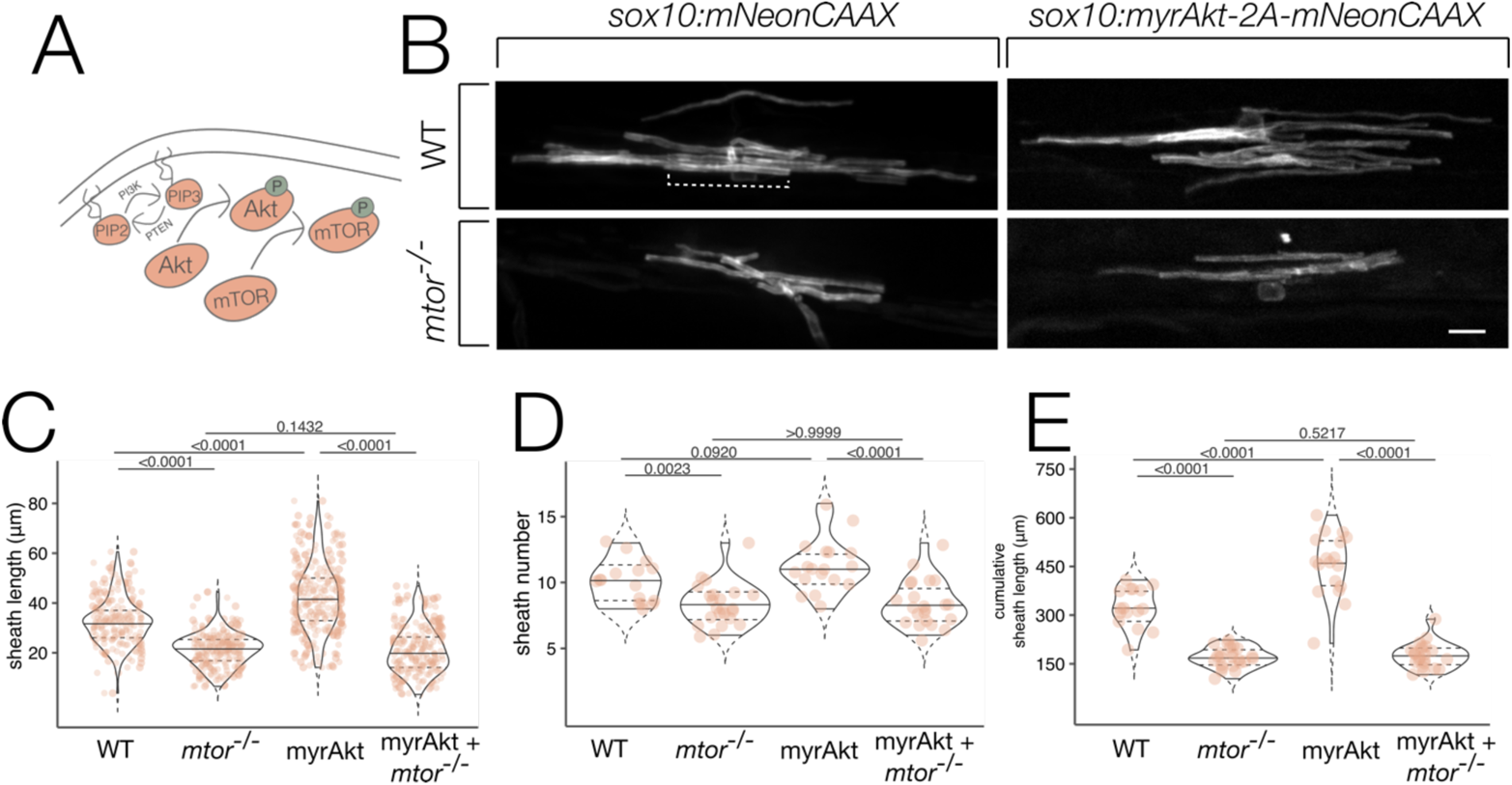
Akt-mTOR pathway activity drives myelin sheath length. (A) Schematic representation of the Akt-mTOR pathway. (B) Representative images of single spinal cord oligodendrocytes of 5 dpf wild-type (WT) or *mtor* mutant larvae with or without expression of myrAkt. Dashed line highlights an individual myelin sheath. Scale bar, 10µm. Quantification of average sheath length (C), sheath number per cell (D), and cumulative myelin sheath length (E). Violin plot solid lines represent median and dashed lines represent 25^th^ and 75^th^ percentiles. Overall significance was assessed using the Kruskal-Wallis test: sheath length, p < 1×10^−12^, h = 362.3; sheath number, p = 3.98×10^−6^, h = 27.81; cumulative sheath length, p = 2×10^−12^, h = 57.40. Individual data points are represented by orange dots. P values on figure are pairwise Mann-Whitney tests with Bonferroni-Holm correction for multiple comparisons. Sample sizes: WT - 16 fish, 16 cells, 161 sheaths; *mtor^−/−^* - 23 fish, 23 cells, 206 sheaths; myrAkt - 18 fish, 18 cells, 197 sheaths; myrAkt + *mtor^−/−^* - 22 fish, 22 cells, 184 sheaths.

In mice, oligodendrocytes programmed to express a constitutively active form of Akt expressed elevated levels of myelin mRNA and protein and produced significantly thicker myelin sheaths (Flores et al., 2008). To test whether elevated Akt signaling also increases myelin sheath length and number, we drove oligodendrocyte-specific expression of a constitutively active Akt1 allele. This allele, hereafter referred to as myrAkt, has an N-terminal myristoylation signal as well as threonine 308 and serine 473, which are the phosphorylation sites that result in full activation, mutated to aspartic acid (Aoki et al., 1998). To identify cells expressing this allele, we appended the viral P2A sequence followed by mNeonCAAX to generate the construct *sox10:myrAkt-P2A-mNeonCAAX,* which drives expression of both myrAkt and mNeonCAAX in oligodendrocyte lineage cells from the same construct. We injected *sox10:myrAkt-P2A-mNeonCAAX* into newly fertilized wild-type eggs and at 5 dpf imaged mNeonCAAX^+^ spinal cord oligodendrocytes for ensheathment analysis. On average, myelin sheaths generated by cells expressing myrAkt were ~30% longer than those made by wild-type controls (Figure 1C). Additionally, cells expressing myrAkt had slightly more sheaths than cells expressing a control, but the difference did not reach statistical significance (Figure 1D). Together, the increased individual myelin sheath length and number resulted in ~40% more myelin generated upon constitutive activation of Akt (Figure 1E). These data show that constitutive Akt activation in wild-type cells autonomously promotes myelin sheath elongation.

In mice, Akt and mTOR function in a common pathway to promote myelin gene and protein expression and myelin sheath thickness (Narayanan et al., 2009). Because Akt has numerous downstream targets, it was important to determine if the myelin sheath length-promoting activity of myrAkt requires mTOR function. To test this, we elevated Akt signaling in oligodendrocytes of *mtor* mutant larvae by injecting *sox10:myrAkt-P2A-mNeonCAAX* into newly fertilized eggs generated from incrosses of *mtor* mutant heterozygotes. If Akt and mTOR function in a common pathway to promote myelin sheath growth, we predicted that driving constitutively active Akt in oligodendrocytes of *mtor* mutant larvae would result in a myelin phenotype indistinguishable from *mtor* mutants alone. By contrast, if Akt does not require mTOR function to promote myelin sheath growth, we predicted driving constitutive Akt activation in oligodendrocytes of *mtor* mutant larvae would increase myelin sheath length. Strikingly, oligodendrocytes in *mtor* mutant larvae that expressed myrAkt produced shorter (Figure 1C) and fewer (Figure 1D) myelin sheaths than wild-type controls. These cells produced an average of ~50% less cumulative myelin sheath lengths than cells of wild-type larvae (Figure 1E). We then compared the myelin profiles of oligodendrocytes of *mtor* mutant larvae with those of *mtor* mutant larvae that also expressed constitutively active Akt and found the average length, number, and cumulative amount of myelin produced per cell were indistinguishable (Figure 1C-E). From this, we conclude that Akt requires mTOR function to promote myelin sheath length, a result that is comparable to the requirement in myelin sheath thickness (Narayanan et al., 2009).

### mTOR pathway activity promotes myelin sheath stabilization

Changes in myelin sheath length could result from differences in growth rate, stability, or both. To understand which of these parameters was altered upon mTOR pathway loss-of-function, we performed time-lapse imaging of actively myelinating oligodendrocytes beginning at 72 hours post fertilization, when myelin sheath growth is highly dynamic. We collected three dimensional images of wild-type transgenic *sox10:mRFP* and *mtor* mutant larvae transiently expressing *sox10:mScarletCAAX* every 15 minutes for 4 hours (Figure 2A). Myelin sheath length was quantified at each timepoint during the imaging period. To understand the net change in myelin sheath length during the imaging period, we subtracted the length of a given sheath in the first frame from its length in the last frame. We found that on average myelin sheaths in wild-type larvae grew longer during the imaging period, whereas sheaths of *mtor* mutant larvae on average became shorter (Figure 2B). We calculated the mean change in sheath length at each time point by subtracting the length in the initial frame and analyzed those as a function of time. On average, sheaths in *mtor* mutant larvae displayed a steady net negative displacement during the imaging period compared to the net positive displacement of sheaths in wild-type larvae (Figure 2C). Together, these data show that mTOR signaling is required to drive myelin sheath growth during development.

**Figure 2.**
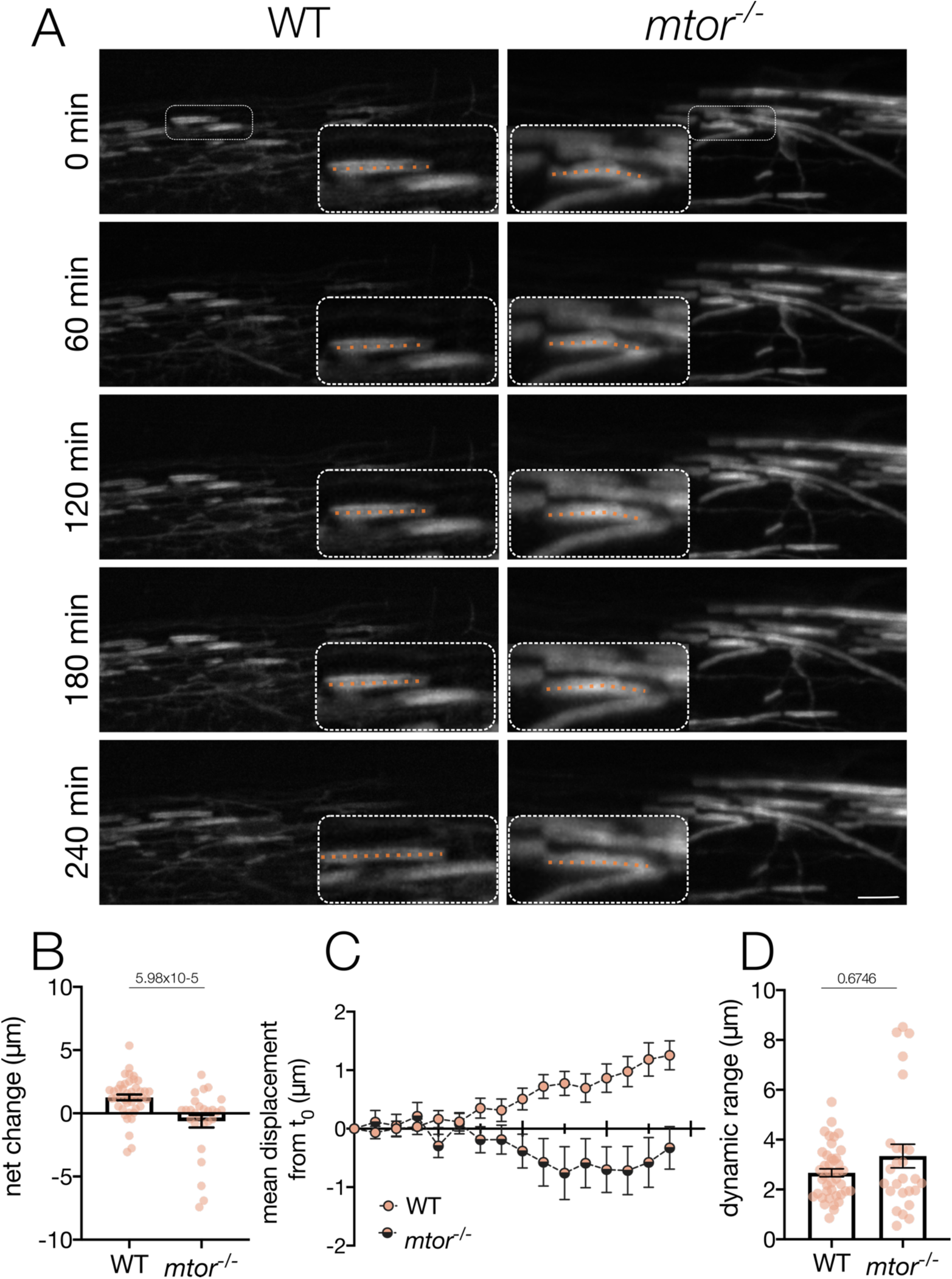
mTOR pathway activity promotes myelin sheath stabilization. (A) Images of myelin sheaths in the spinal cord of 72 hours post fertilization wild-type (WT) or *mtor* mutant larvae at 60 minute intervals during the imaging period. Inset, dashed line highlights an individual myelin sheath throughout the time course. Scale bar, 10µm. (B) Average net change (length at 240 minutes – length at 0 minutes) of sheaths in WT and *mtor* mutant larvae. (C) Mean change in sheath length from initial length as a function of time. Mann-Whitney, p = 5.98×10^−5^, U = 258. (D) Dynamic range (maximum length – minimum length) of sheaths in WT and *mtor* mutant larvae. Mann-Whitney, p = 0.6746, U = 512. N = WT - 8 animals, 42 sheaths; *mtor^−.−^* - 7 animals, 28 sheaths.

Why did myelin sheaths in *mtor* mutant larvae not show a net growth during development? Did these sheaths not grow at all or did they grow but were not stabilized? To distinguish between these possibilities, we calculated the total amount of space an individual myelin sheath occupied during the imaging period. This measure, which we referred to as dynamic range, is defined by the maximum length of a given sheath subtracted by the minimum length of that sheath. There was no difference in the average dynamic range of myelin sheaths in wild-type and *mtor* mutant larvae (Figure 2D), which shows that in the absence of mTOR signaling myelin sheaths are capable of growth but are not stabilized.

### Oligodendrocytes localize mTOR-regulated translational machinery to myelin sheaths

The localization of mRNAs (Ainger et al., 1993; Thakurela et al., 2016; Yergert et al., 2021), ribosomes (Lunn et al., 1997), and RNA binding proteins (Doll et al., 2020) in distal myelin sheaths support a model where local protein translation promotes myelination. We therefore investigated the possibility that translation factors that are regulated by mTOR signaling are localized to nascent myelin sheaths. To test this possibility, we generated a C-terminally tagged human 4E-BP1 fusion protein, an mTOR-dependent translational regulator, and expressed it specifically in oligodendrocytes using the Tol2-transgenesis system and *sox10* regulatory DNA. Imaging of oligodendrocytes at 4 dpf revealed that oligodendrocytes localized 4E-BP1-eGFP to myelin sheaths (Figure 3A). N-terminally tagged fusion proteins were similarly localized, indicating that sheath localization is not a consequence of the placement of the eGFP fusion. The localization of 4E-BP1-eGFP was similar to localization of F-actin that defines the leading edge of myelin sheaths (Figure 3A’). These observations therefore raise the possibility that translation regulated by 4E-BP1 downstream of Akt-mTOR signaling occurs at the leading edge of myelin sheaths.

**Figure 3.**
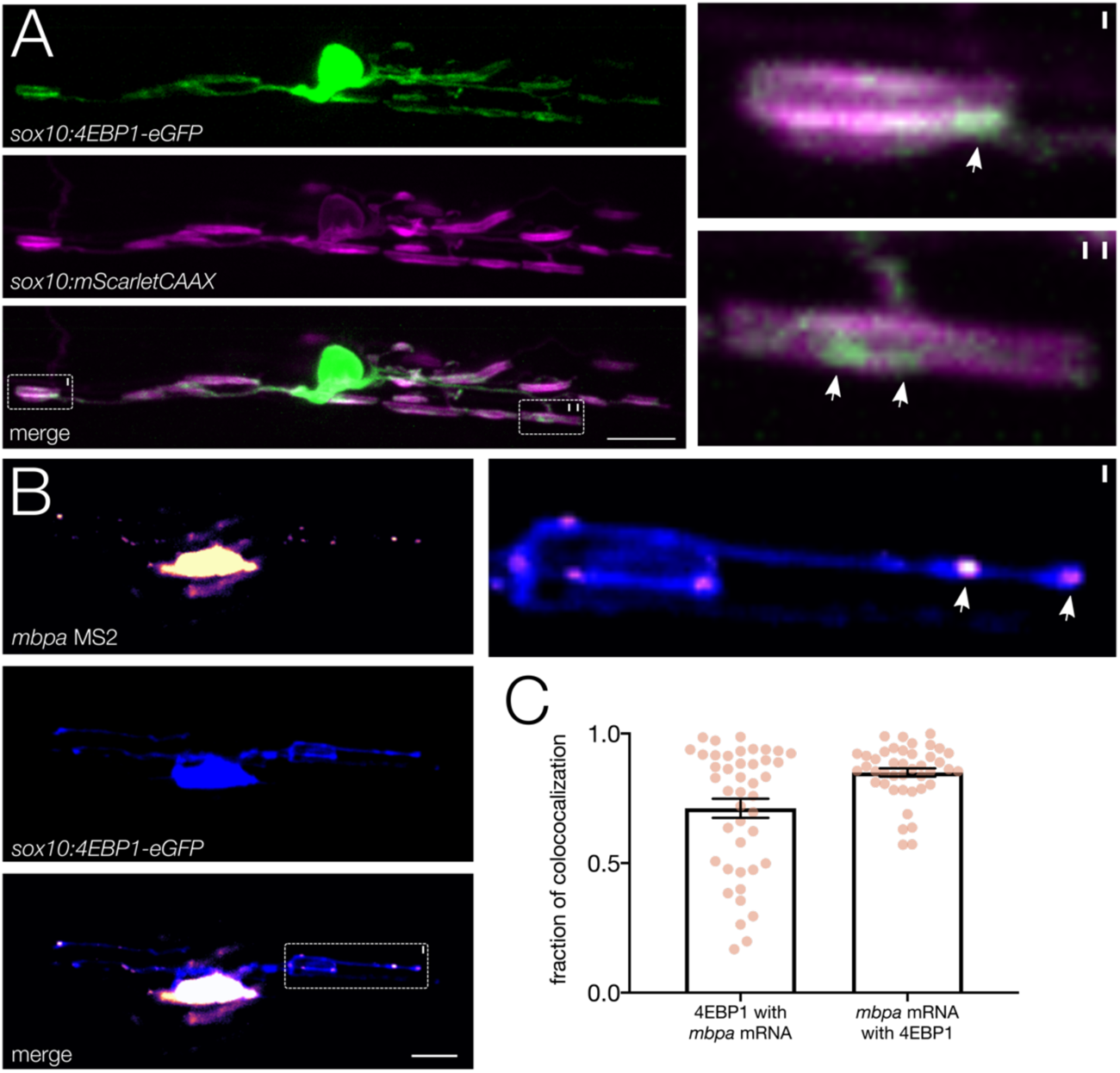
Oligodendrocytes colocalize translational regulators and mRNA in myelin. (A) Image of single 4 dpf spinal cord oligodendrocyte co-expressing *sox10:mScarletCAAX* to label membrane and *sox10:4EBP1-eGFP*. 4EBP1-eGFP signal observed in myelin sheaths and localized to the distal ends (inset, arrows). Scale bar, 10µm. (B) Colocalization of 4EBP1-eGFP and *mbpa* mRNA MS2 reporter in a spinal cord oligodendrocyte. 4EBP1-eGFP and *mbpa* colocalize to discrete puncta in the myelin sheath (inset, arrows). (C) Quantification of 4EBP1-eGFP and *mbpa* mRNA colocalization in individual myelin sheaths. Sample size: 7 animals, 7 cells, 43 sheaths.

For Akt-mTOR signaling to promote localized translation, we predicted that Akt-mTOR-dependent translational regulators would co-localize with myelin localized mRNAs. To investigate this possibility, we used fluorescent reporters to test if 4E-BP1 colocalized with *mbpa* mRNA, which encodes myelin basic protein, in vivo using the MS2-MCP RNA visualization system. The MS2-MCP system consists of an mRNA containing 24 MS2 binding sites (24xMBS), which form repetitive stem loops, and a fluorescently tagged MS2 coat protein (MCP), which specifically binds the 24xMBS stem loops to visualize mRNA via fluorescent reporter (Bertrand et al., 1998). Our lab has previously used this system to reveal localization of RNA encoded by *mbpa,* a zebrafish orthologue of *Mbp,* to myelin sheaths (Yergert et al., 2021). We co-expressed 4E-BP1-eGFP and *mbpa* MS2-mScarlet in oligodendrocytes using the Tol2-transgenesis system and imaged eGFP/mScarlet^+^ spinal cord oligodendrocytes at 4 dpf (Figure 3B). Expression of 4E-BP1-eGFP and *mbpa* MS2 was highly colocalized in myelin sheaths (Figure 3B, inset). We performed colocalization analysis using JACoP (Just Another Colocalization Plugin) and calculated Mander’s coefficient to quantify the amount of 4E-BP1-eGFP colocalized with *mbpa* mRNA and vice versa. We found that ~85% of *mbpa* mRNA colocalized with 4E-BP1-eGFP in myelin sheaths (Figure 3C), while 4E-BP1-eGFP colocalized with *mbpa* mRNA ~70% of the time (Figure 3C). Additionally, the percent of 4E-BP1-eGFP that colocalized with *mbpa* mRNA was highly variable, raising the possibility that 4E-BP1 regulates the translation of other myelin-resident mRNAs.

### Identification of putative translational targets in myelin

What myelin-resident mRNAs might be targets of mTOR-regulated translation? To address this question, we used bioinformatics to identify putative myelin-resident translational targets. Loss of 4E-BPs in cultured cells was sufficient to render translation of mRNAs containing terminal oligopyrimidine (TOP) or TOP-like motifs in their 5’ untranslated region (5’ UTR) resistant to inhibition by the mTOR inhibitor Torin 1 (Thoreen et al., 2012), indicating that mTOR signaling through 4E-BPs promotes the translation of 5’ TOP or TOP-like motif containing transcripts. To identify putative 4E-BP1 translational targets, we used the function Find Individual Motif Occurrences (FIMO) in the package MEME Suite to scan the myelin transcriptome for TOP or TOP-like motifs (Figure 6A). We found that 324 of the 1195 annotated 5’UTRs in the myelin transcriptome contained a TOP-like motif (Figure 4B; Table 4-1), indicating that nearly 25% of mRNAs in the myelin transcriptome could be subject to translational regulation by mTOR-4E-BP1 signaling. Canonically, TOP-motif containing mRNAs encode proteins that are components of translational machinery (Yoshihama et al., 2002; Iadevaia et al., 2008). We identified such proteins in our dataset, including the 40S ribosomal proteins S2 and S18 (Rps2 and Rps18, respectively). Additionally, we identified other transcripts with known TOP motifs, such as vimentin (Meyuhas and Kahan, 2015), together indicating that our bioinformatic analysis accurately identified TOP and TOP-like motif containing mRNAs in the myelin transcriptome. To ask if myelin-resident mRNAs with TOP or TOP-like motifs encode specific functions, we performed gene ontology (GO) analysis. From this, we identified some processes indicative of developmental myelination, such as cell junction, membrane, nervous system development, and cytoplasm (Figure 4C). Similar to previous analysis of myelin-resident transcripts (Yergert et al., 2021), we also identified terms associated with synapses and cell signaling (Figure 4C). These data support the possibility that 4E-BP1 regulated translation of specific myelin-resident mRNAs could promote myelination via synaptogenic-like mechanisms.

**Table 4-1.**
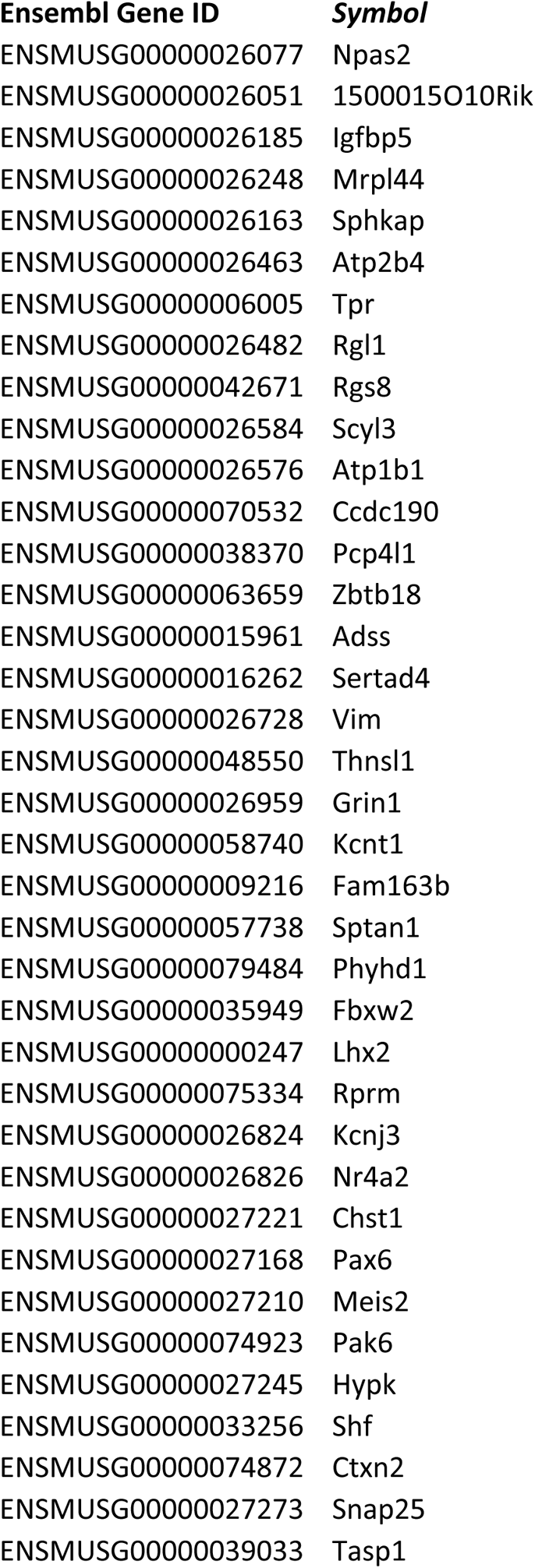
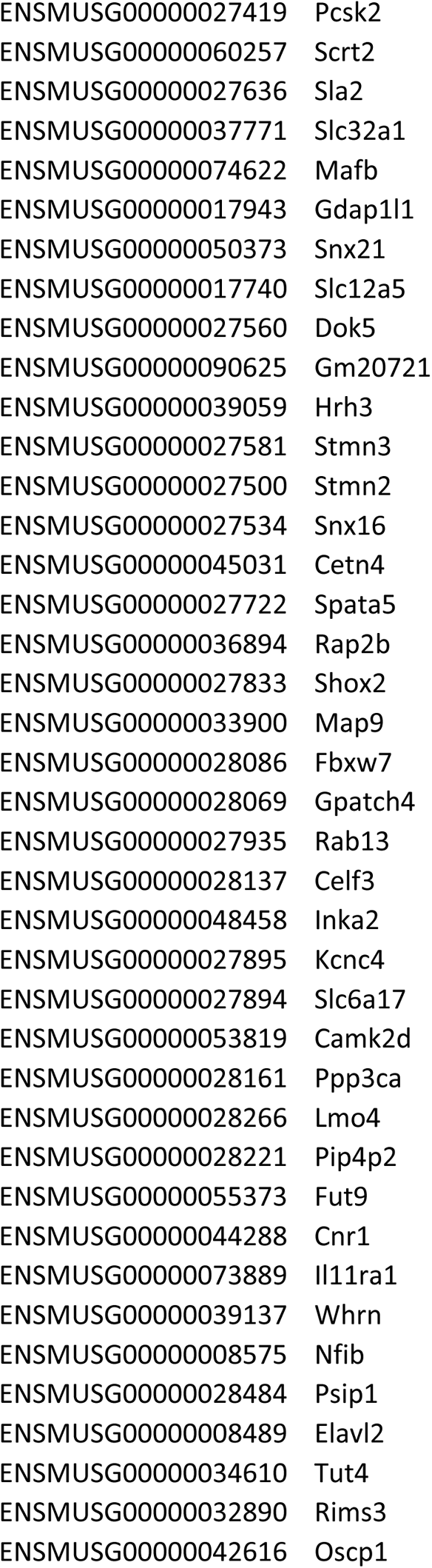
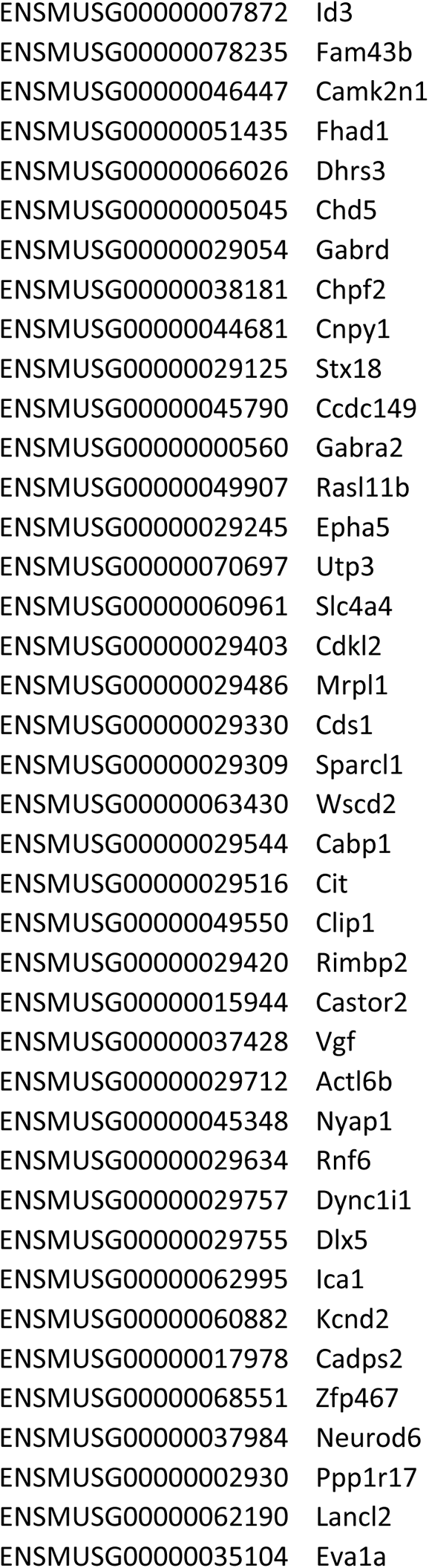
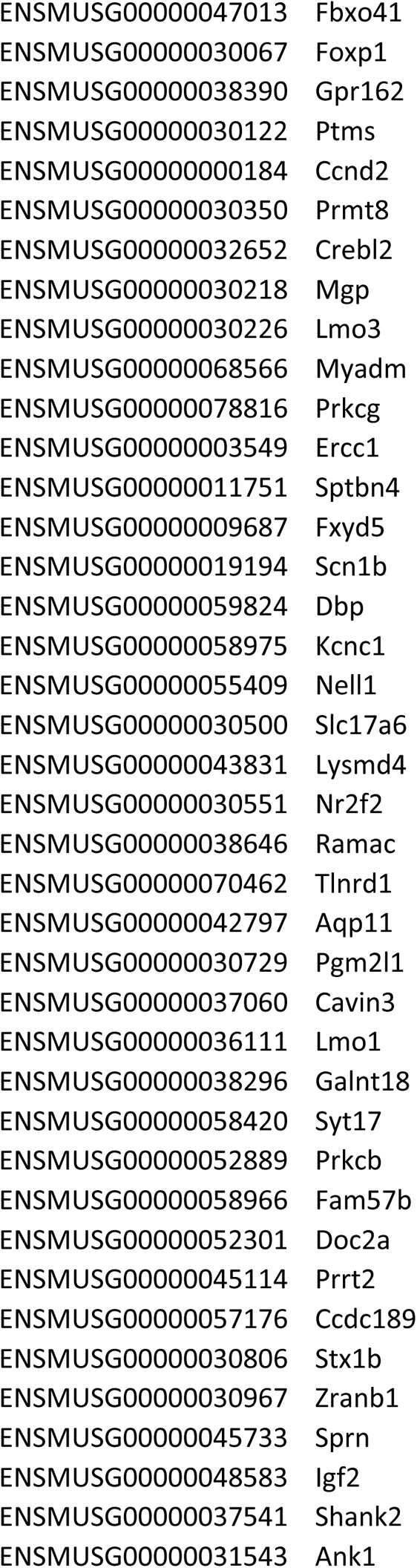
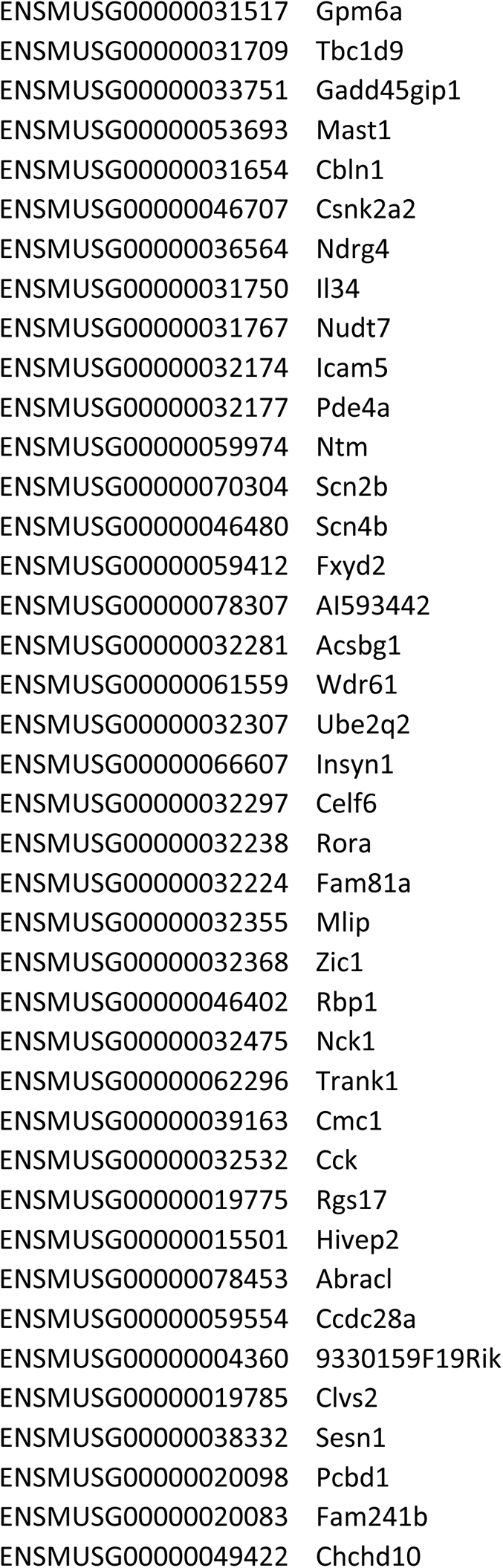
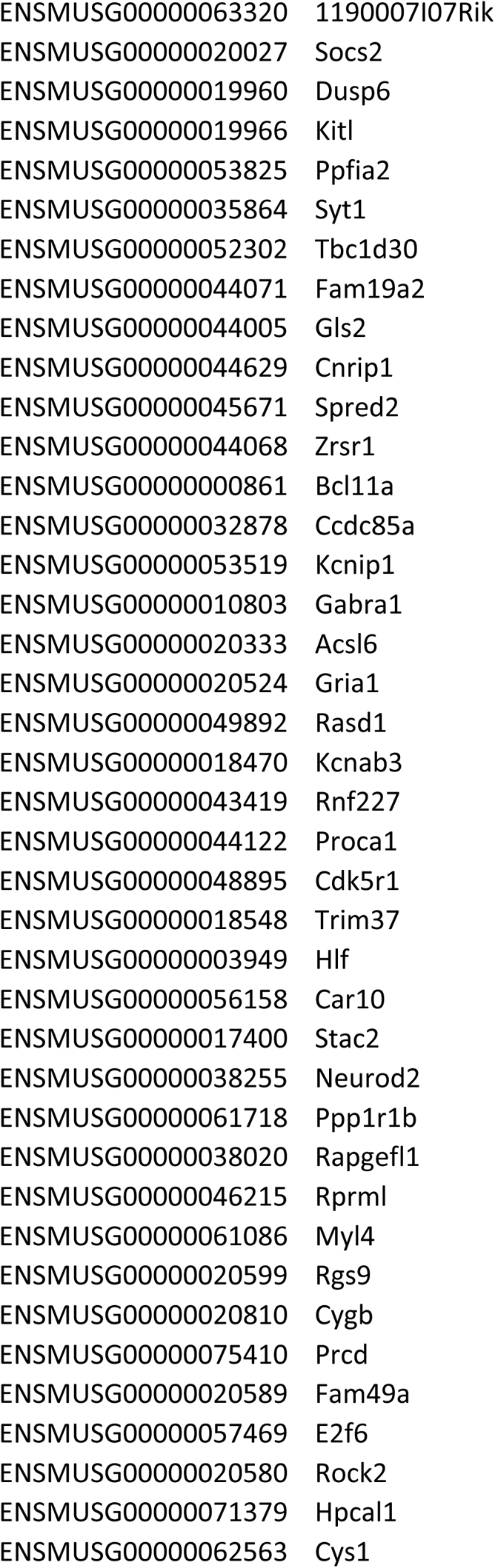
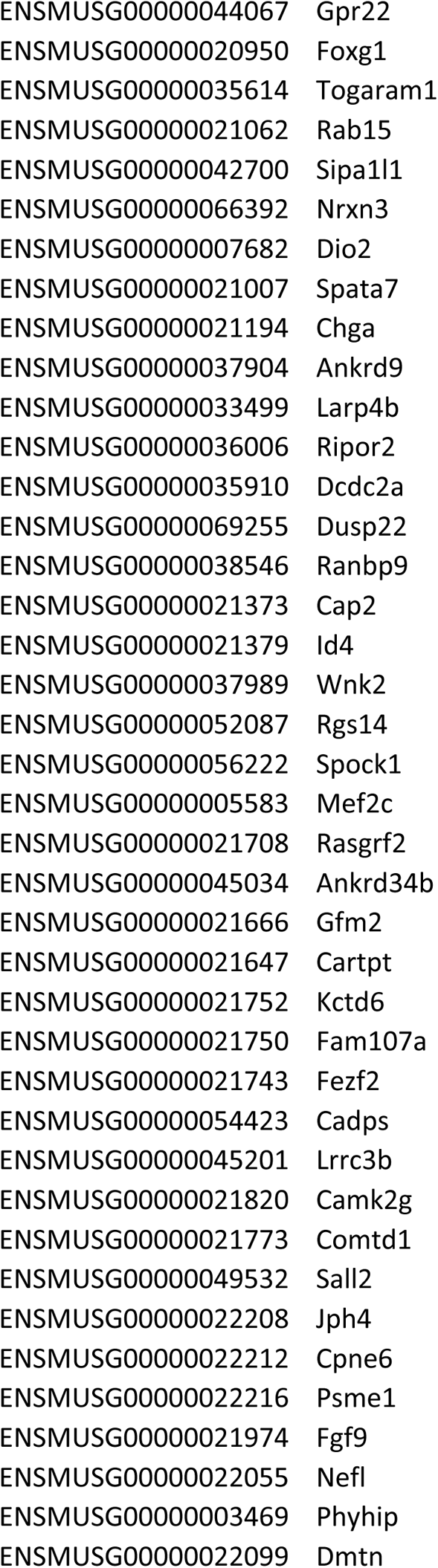
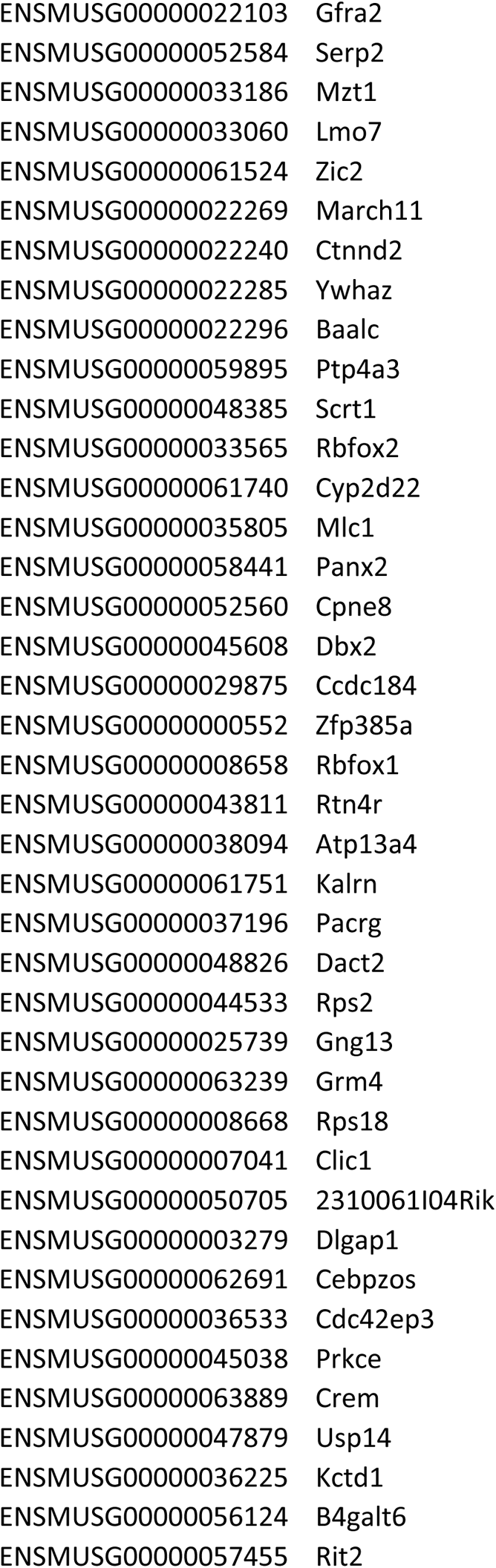
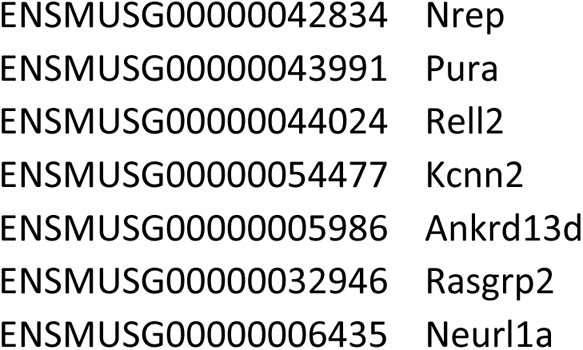
List of genes that are putative 4E-BP1 translational targets in myelin

**Figure 4.**
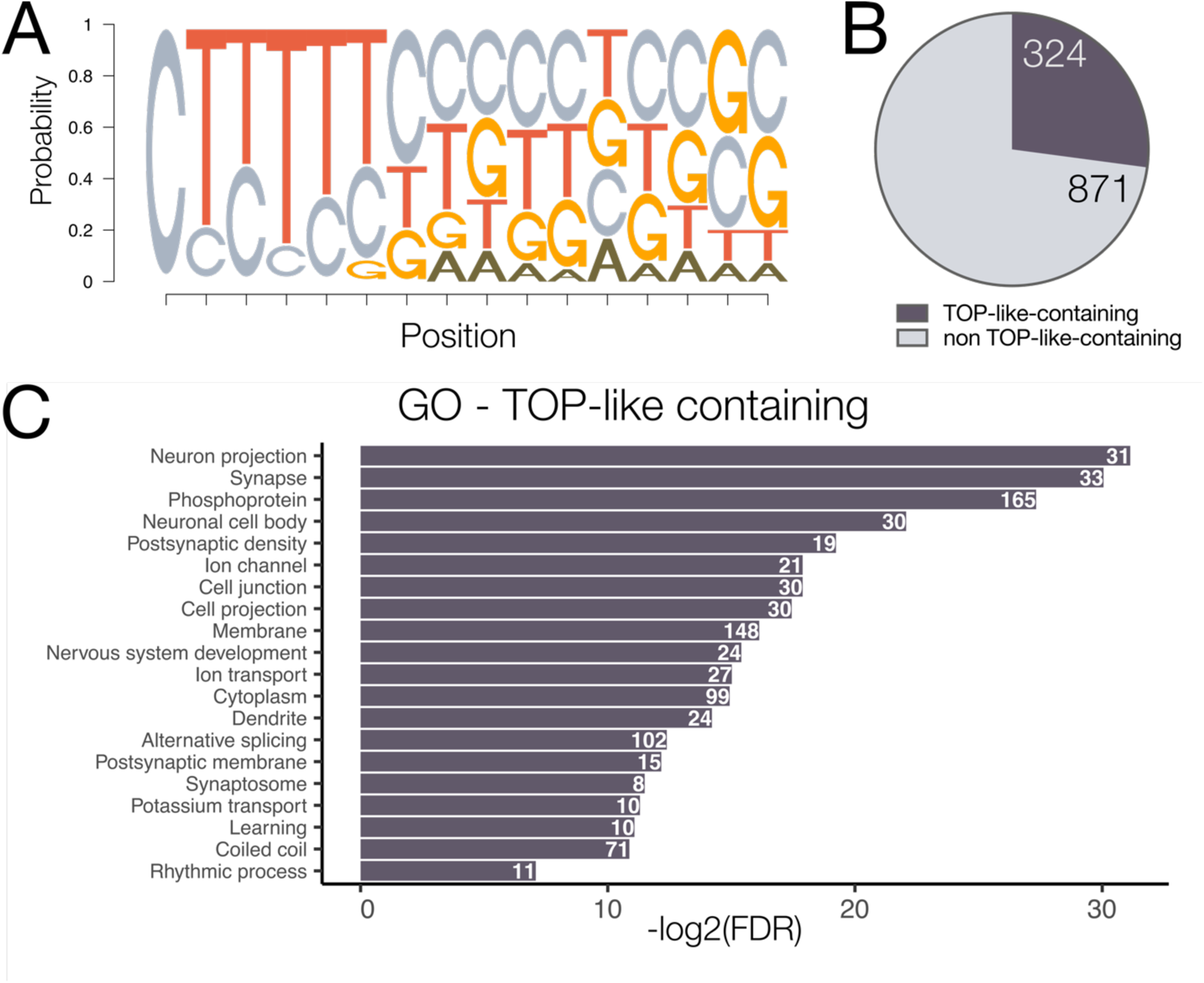
Identification of putative myelin-resident 4E-BP1 translational targets. (A) Schematic representation of the TOP-like motif position weight matrix used in bioinformatic identification of putative translational targets. (B) Percentage of 5’UTRs identified as containing a TOP or TOP-like motif. (C) Gene ontology of transcripts containing identified TOP-like motifs. Top 20 terms shown, ordered most to least significant by −log2 of false discovery rate. Counts correspond to number of genes associated with each term.

We next asked if any myelin-resident mRNAs were putative translational targets of the 4E-BP1 binding partner eIF4E. Studies from eIF4E haploinsufficient mice identified the shared *cis*-regulatory element Cytosine Enriched Regulator of Translation (CERT) present in ~70% of mRNAs that are translational targets of eIF4E (Truitt et al., 2015). To identify transcripts containing a CERT sequence in the myelin transcriptome, we used the FIMO function in MEME suite to search the 5’UTRs of myelin resident transcripts for CERT (Figure 5A). We found that nearly 50% (570/1195) of the annotated 5’UTRs in the myelin transcriptome contain the CERT sequence (Figure 5B; Table 5-1). Similar to 4E-BP1 targets CERT-containing transcripts were enriched for gene ontology terms associated with synapses and intracellular signaling, but also included different categories such as lipoprotein and ion channel (Figure 5C).

**Table 5-1.**
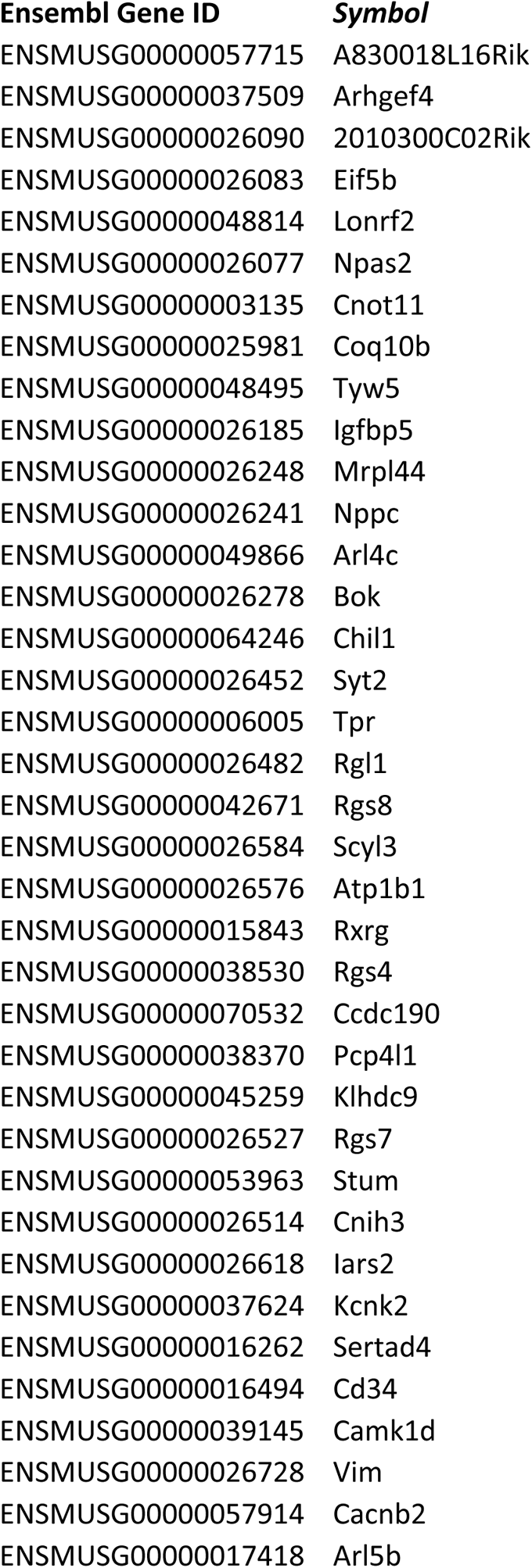
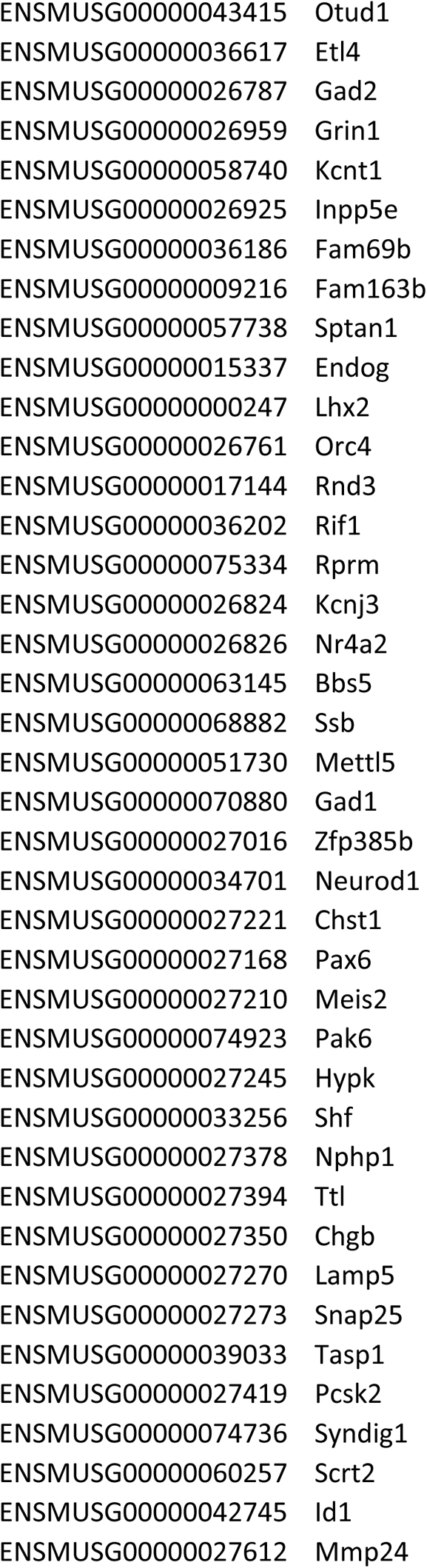
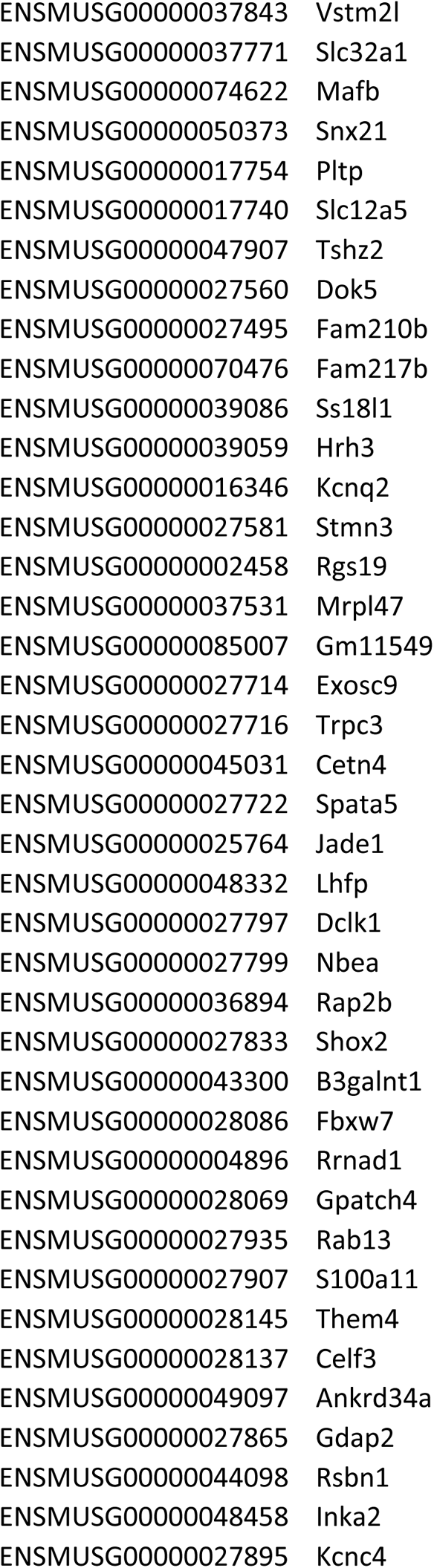
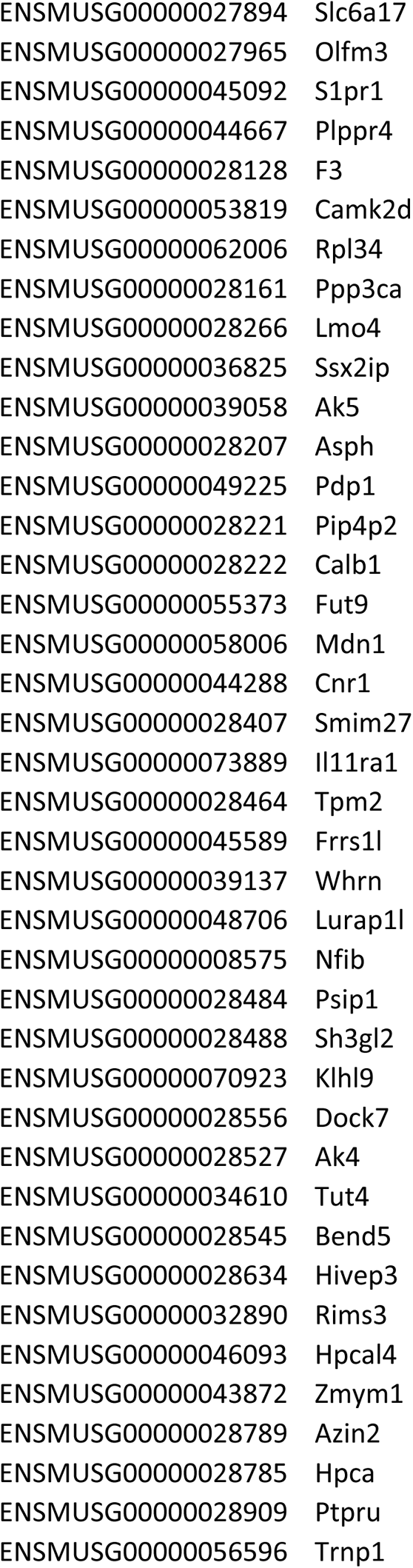
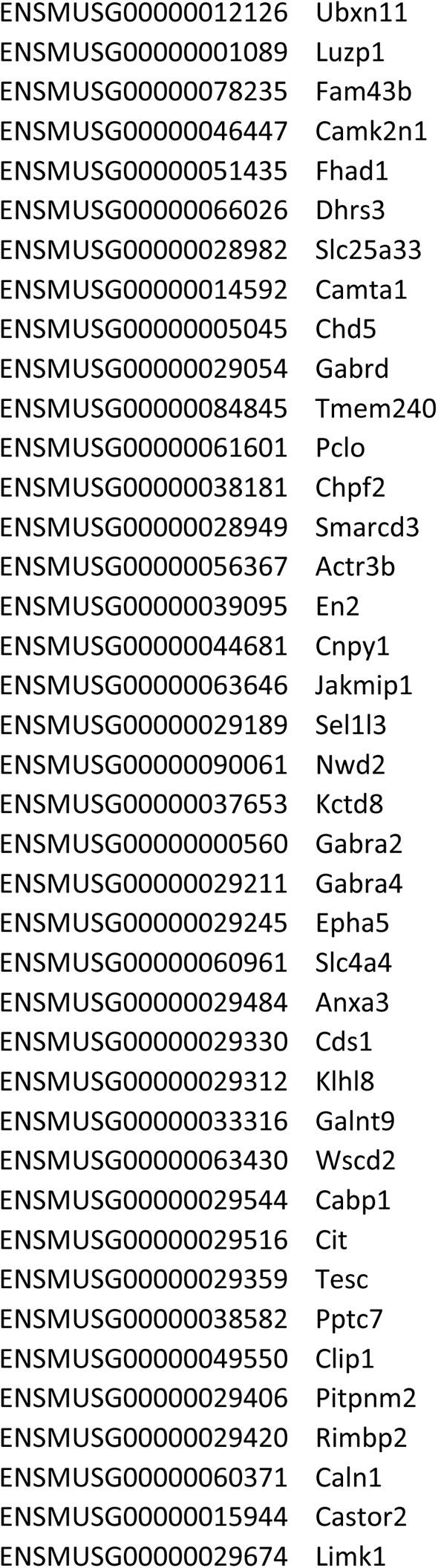
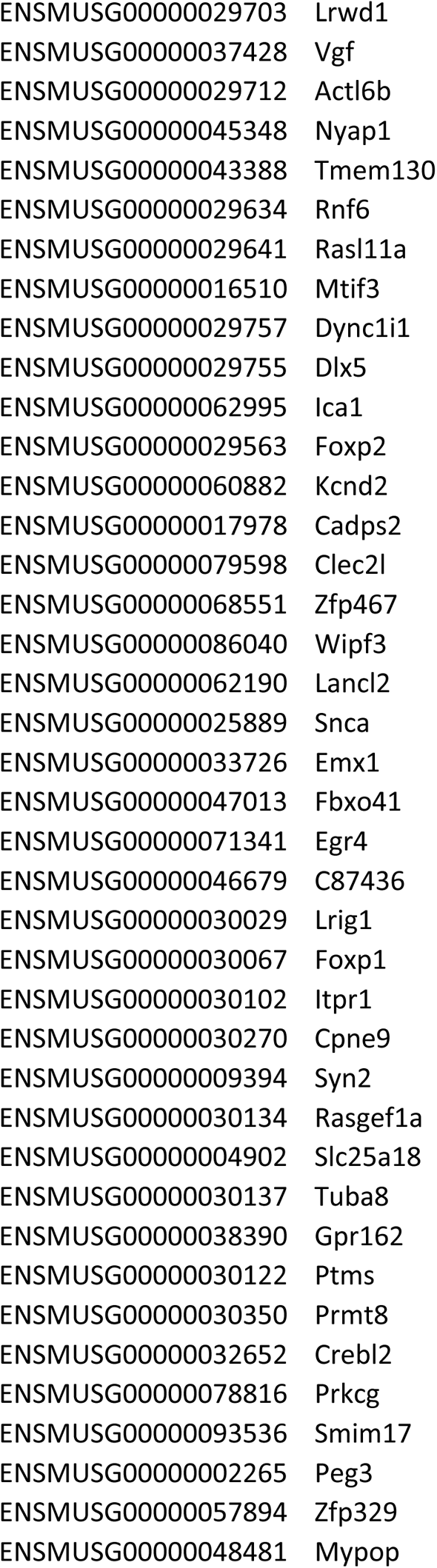
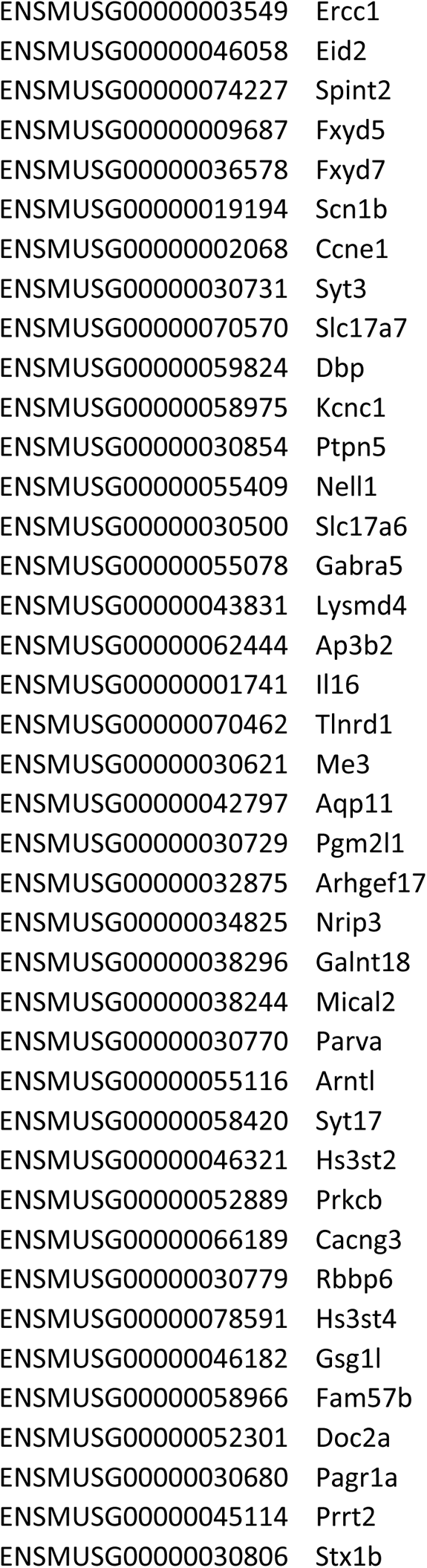
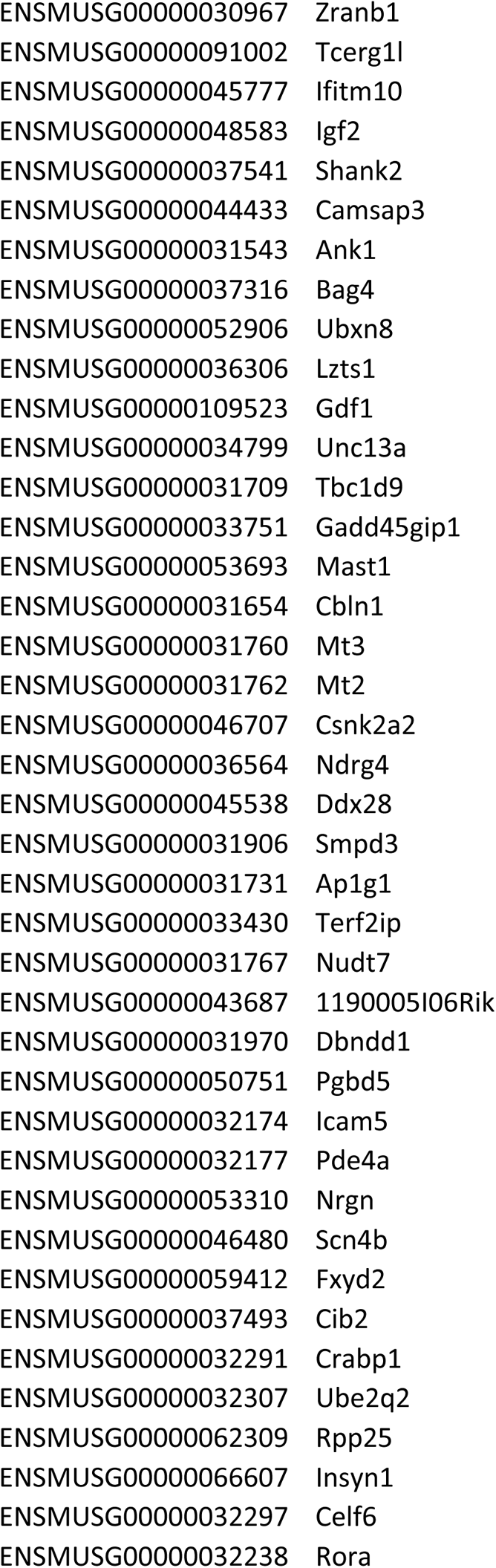
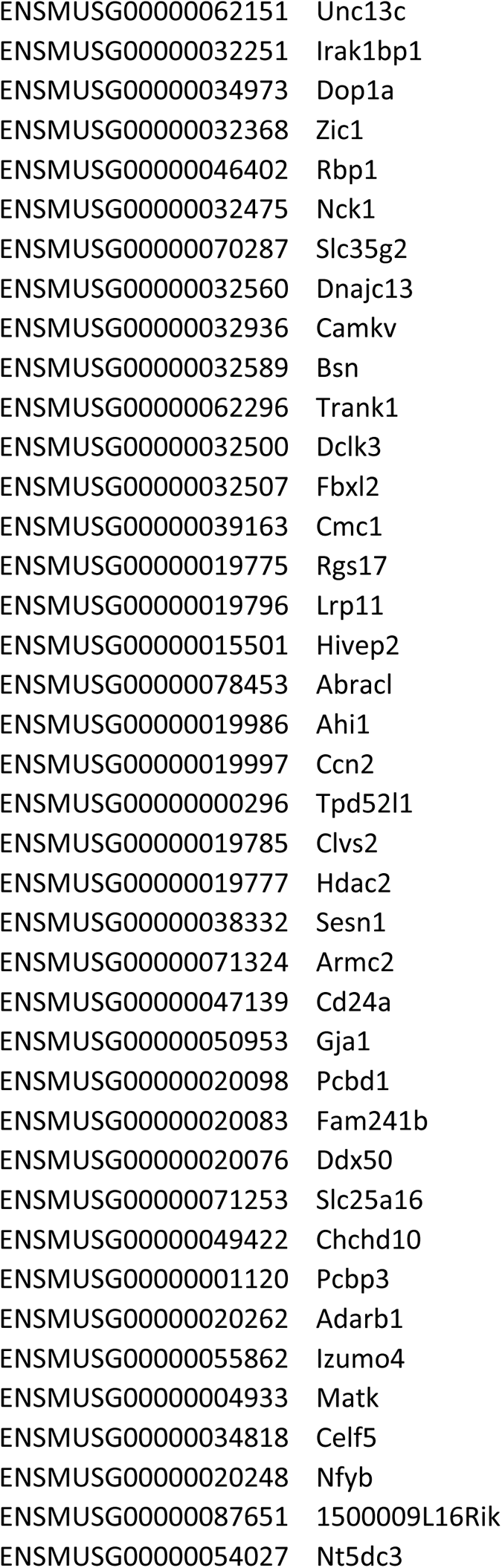
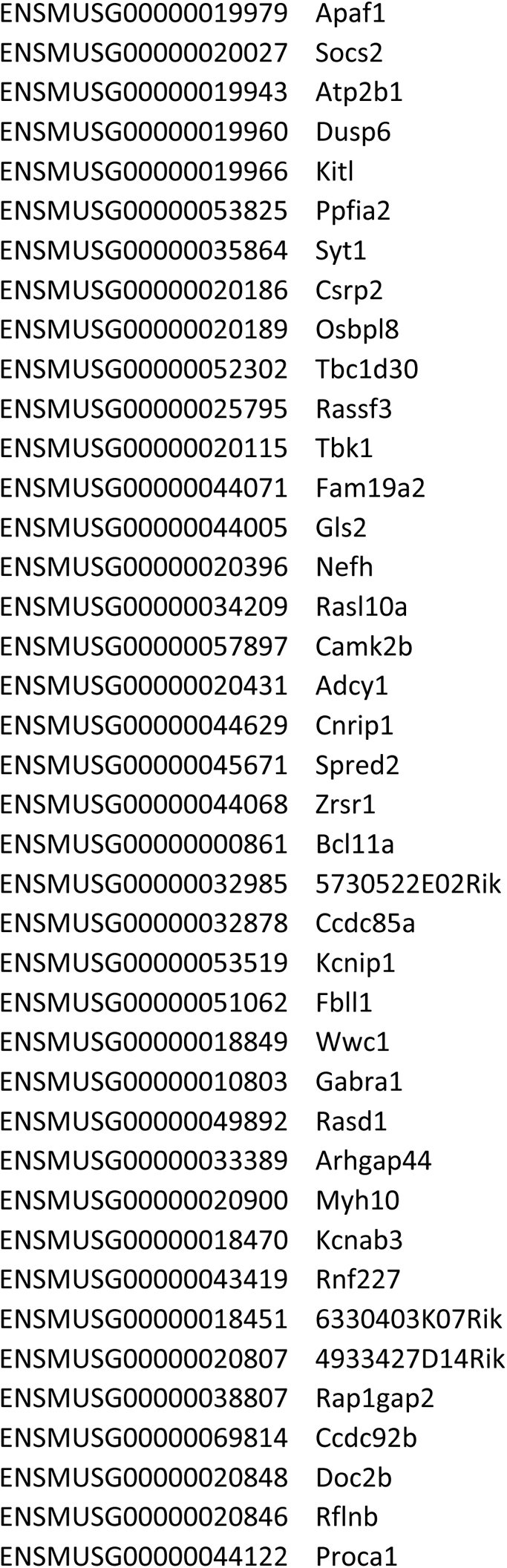
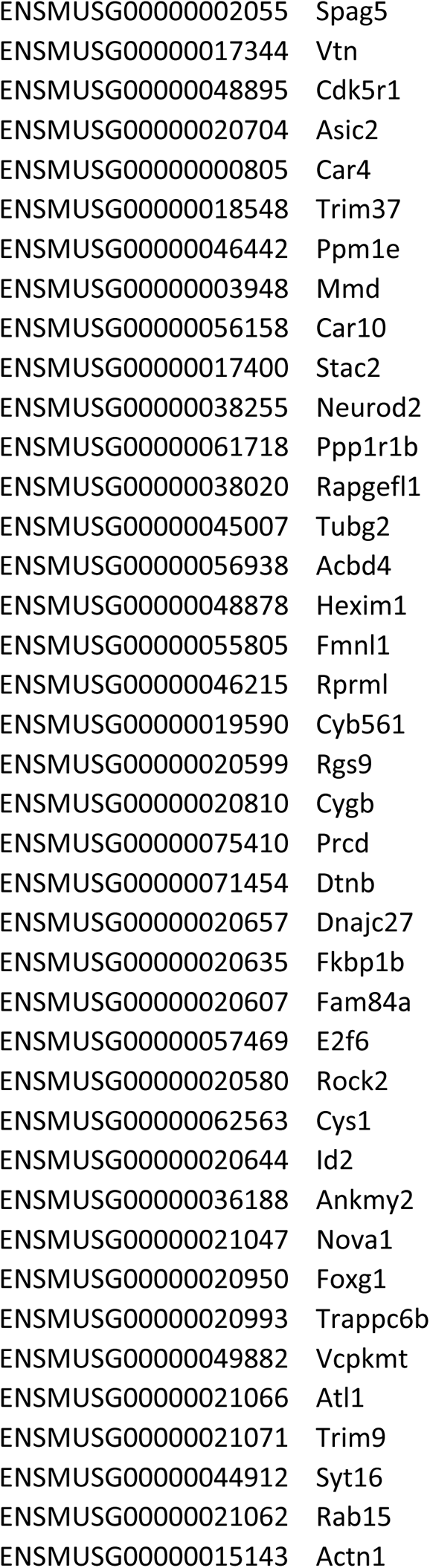
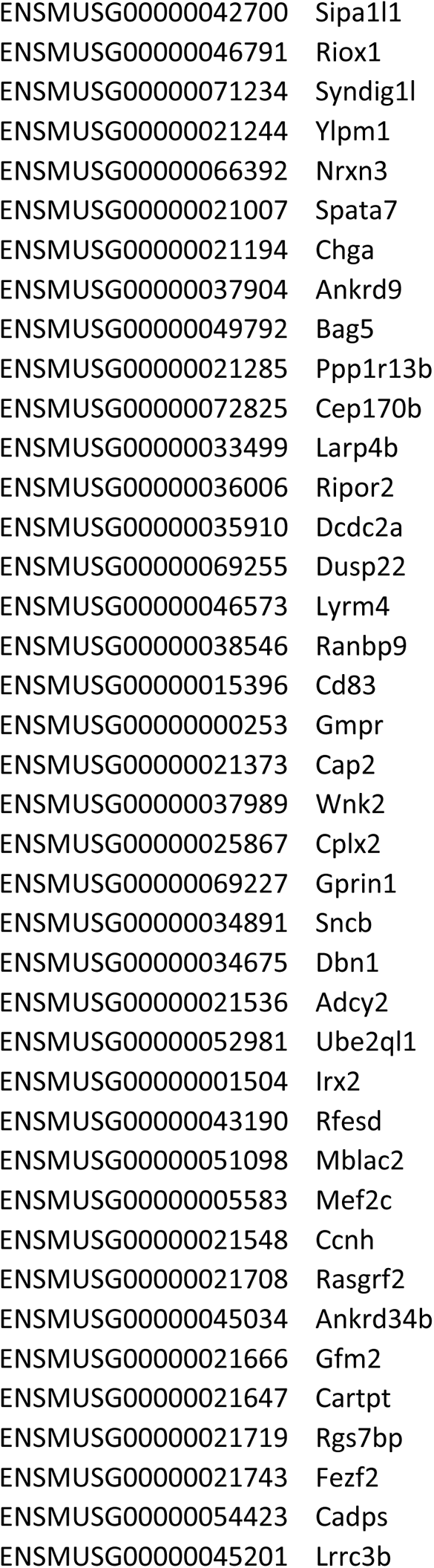
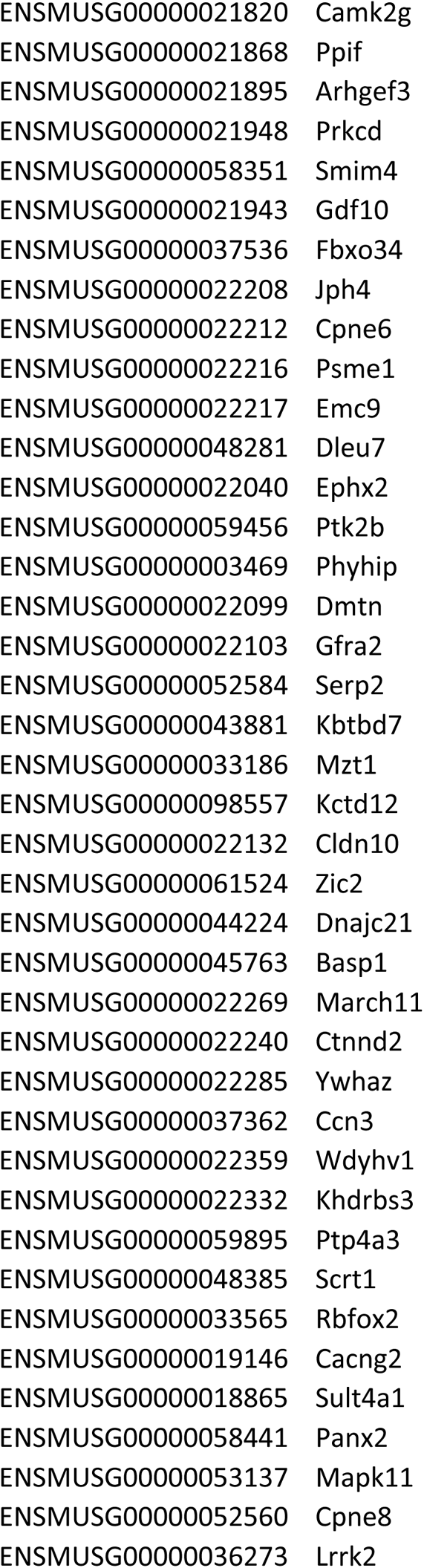
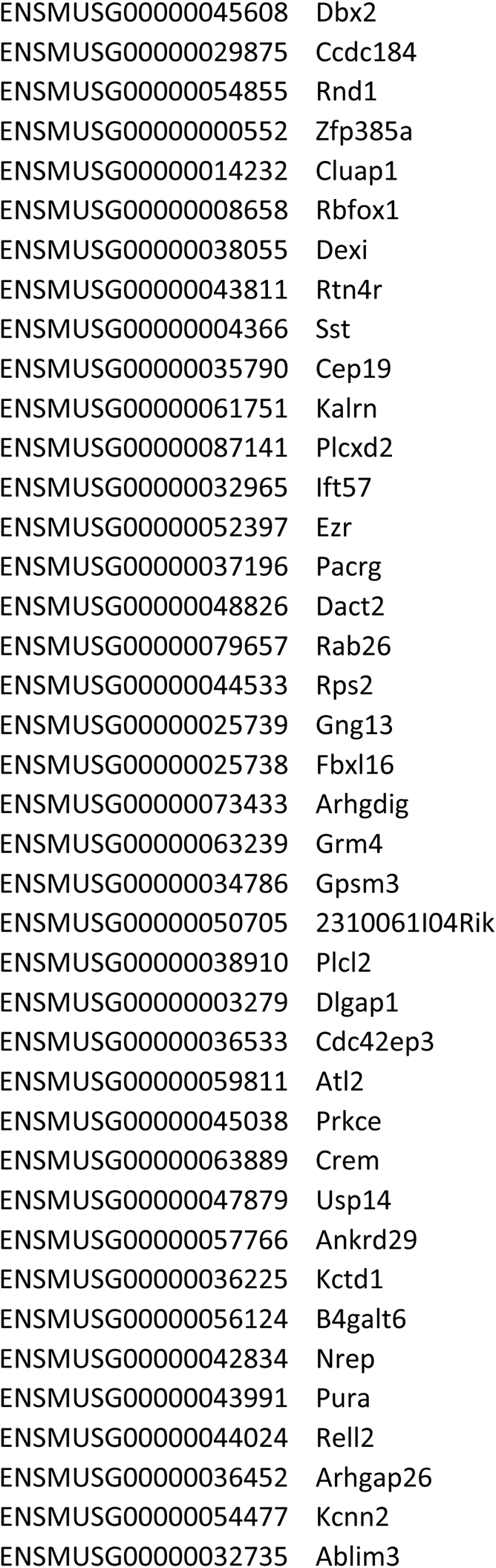
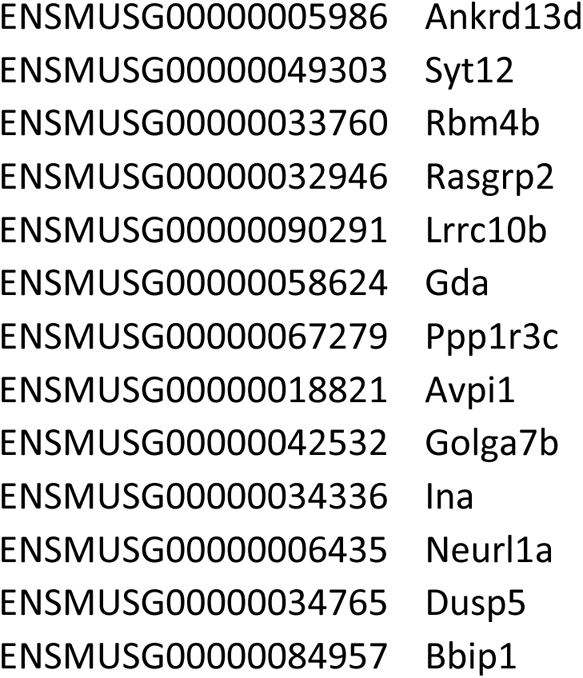
List of genes that are putative eIF4E translational targets in myelin

**Figure 5.**
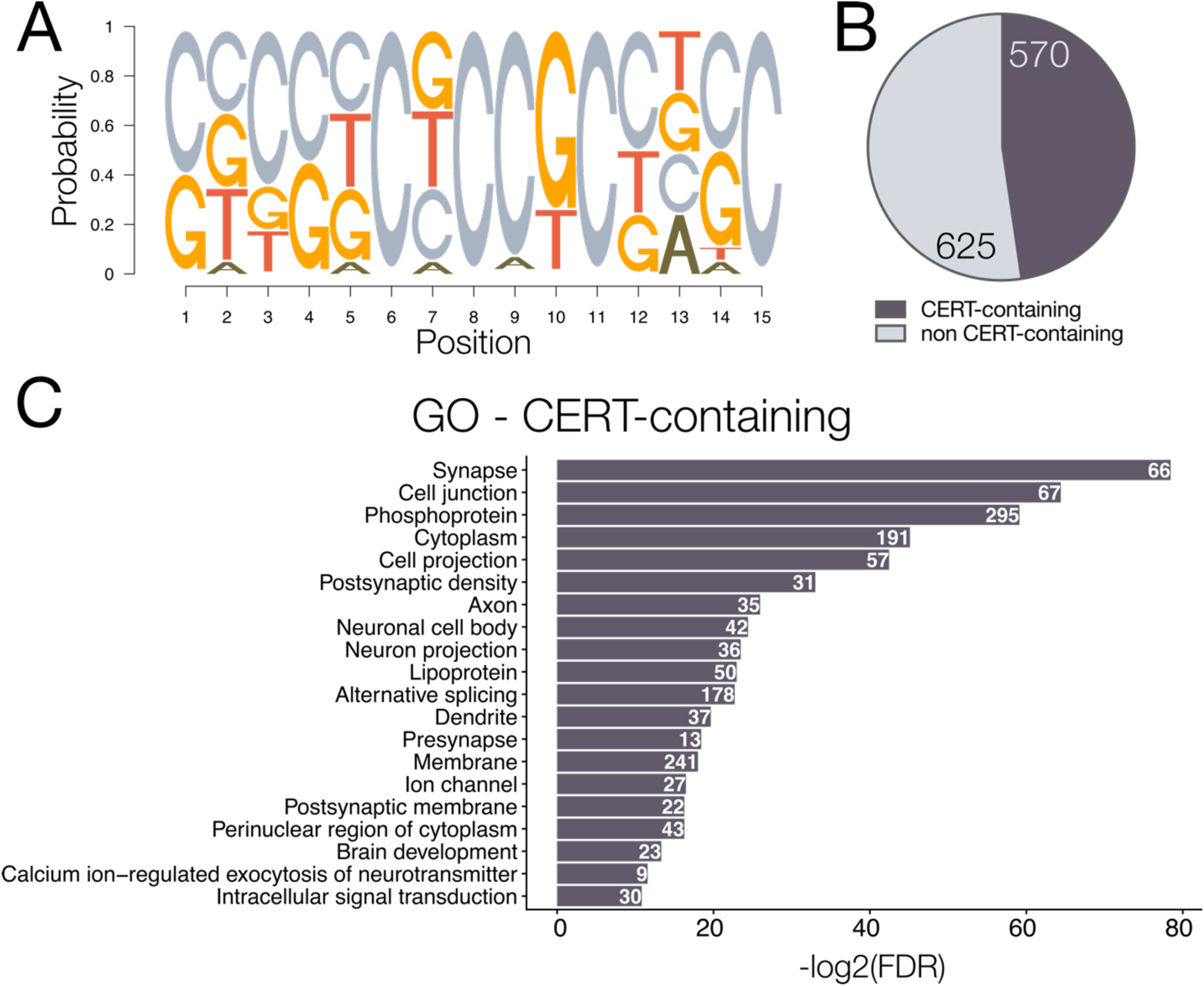
Identification of putative eIF4E myelin-resident translational targets. (A) Schematic representation of CERT position weight matrix used to identify transcripts of interest. (B) Percentage of myelin-resident mRNAs identified as having a CERT motif in the 5’UTR. (C) Gene ontology of transcriptions containing identified CERT motif. Top 20 terms shown, ordered most to least significant by −log2 of false discovery rate. Counts correspond to number of genes associated with each term.

Because both 4E-BP1 and eIF4E regulate translation downstream of mTOR signaling, we defined the transcripts that contain both of these sequences. Of the 1195 annotated 5’UTRs in the myelin transcriptome, 245 possessed both a TOP-like motif and a CERT sequence (~20%) (Figure 6A). Approximately 75% of TOP-like motif containing also contained a CERT sequence (245/324) (Figure 6B; Table 6-1), whereas only ~40% of CERT-containing transcripts also contain a TOP-like motif (245/570) (Figure 6B). Myelin-resident transcripts containing both a TOP-like motif and CERT sequence represented GO terms associated with myelin development such as cell junction, membrane, and nervous system development, but also represented distinct categories such as those associated with potassium and neurotransmitter transport (Figure 6C). Together, the similarities and differences among putative 4E-BP1 and eIF4E translational targets supports the possibility that translation regulated by these two proteins have both overlapping and distinct functions in myelin development.

**Table 6-1.**
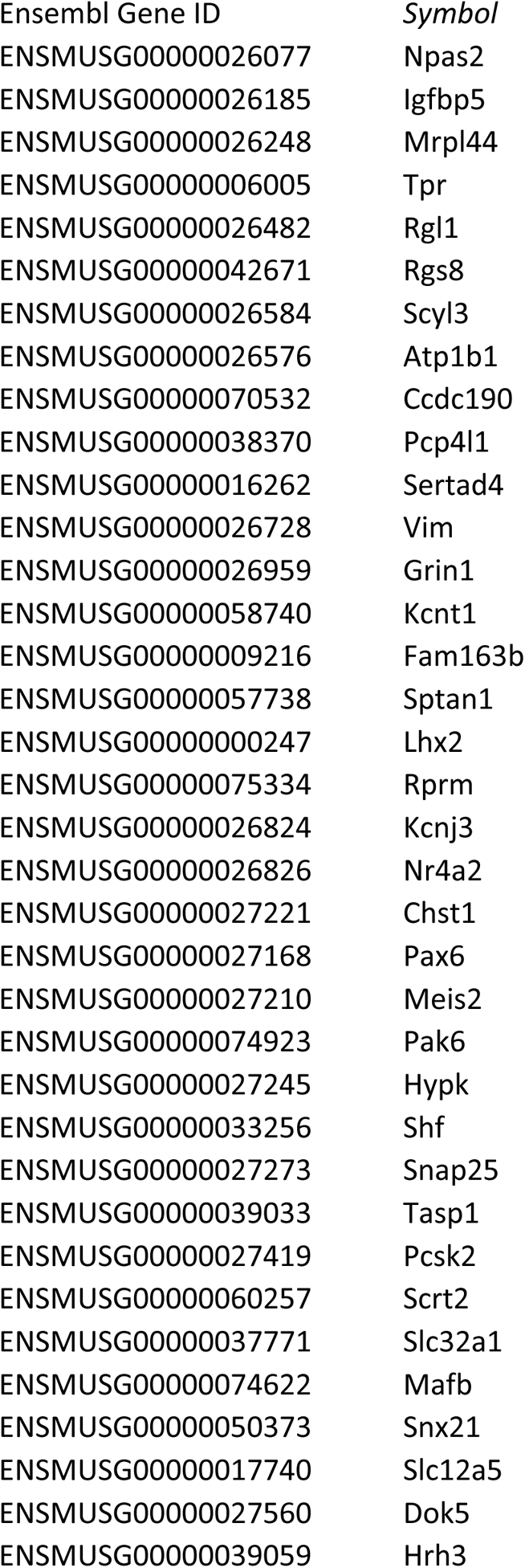
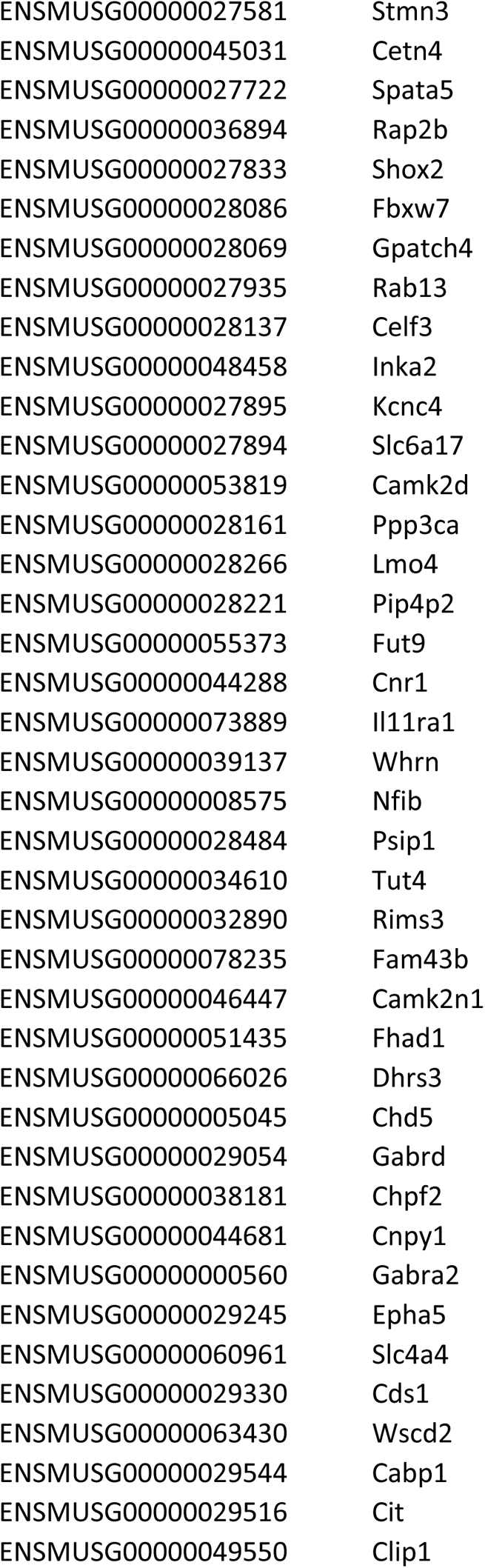
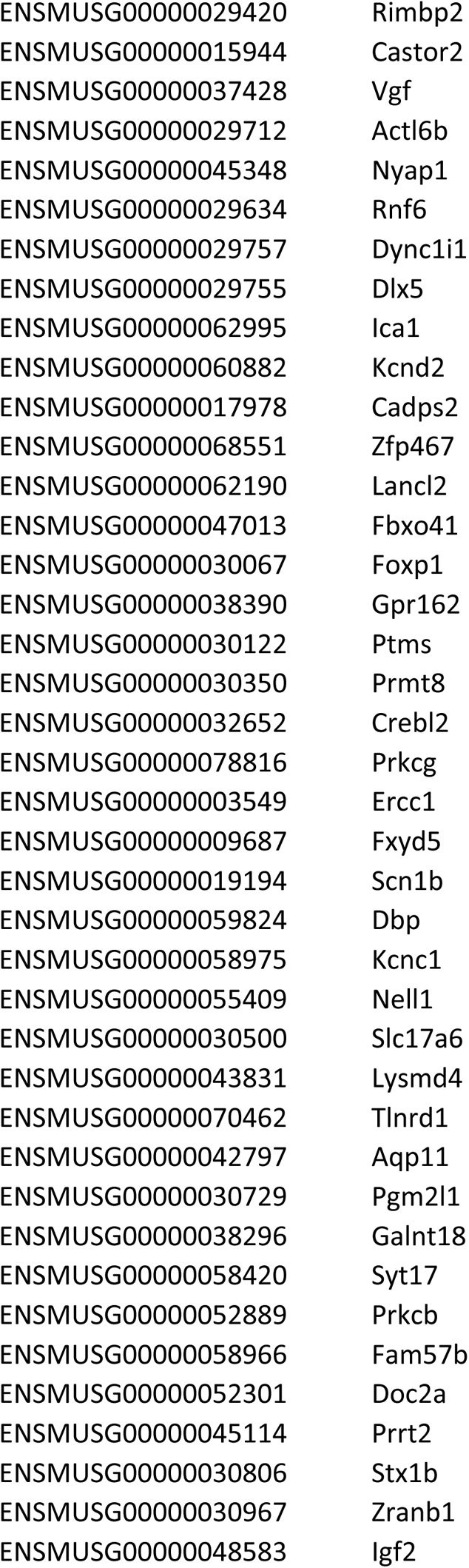
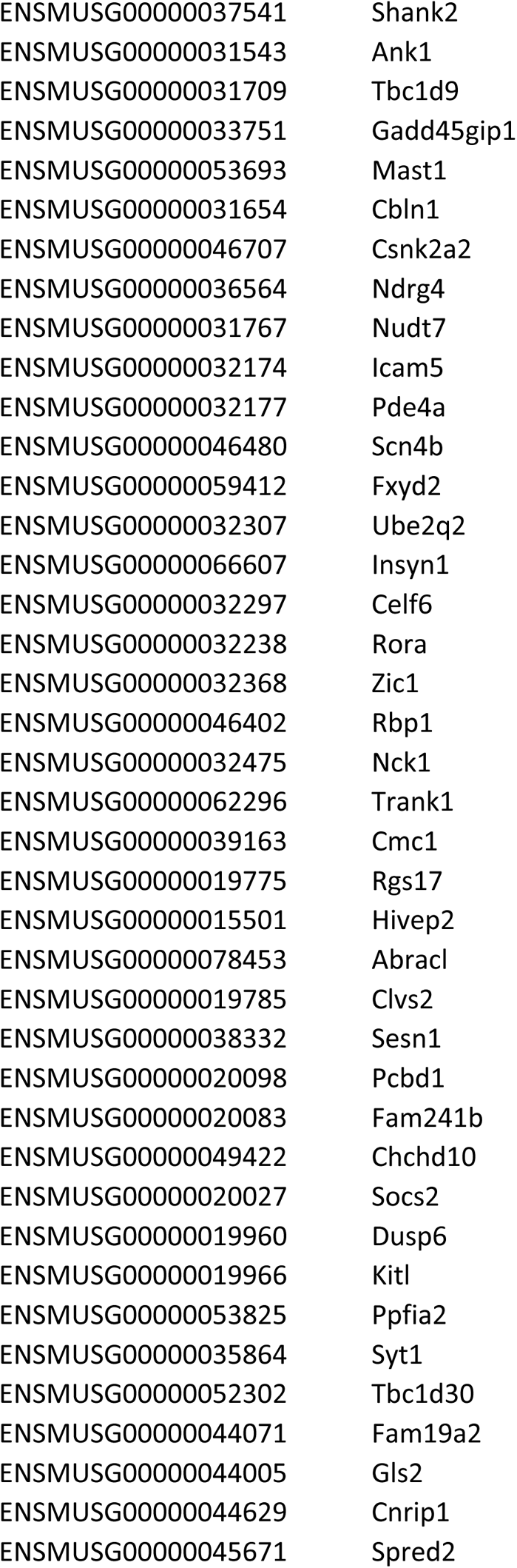
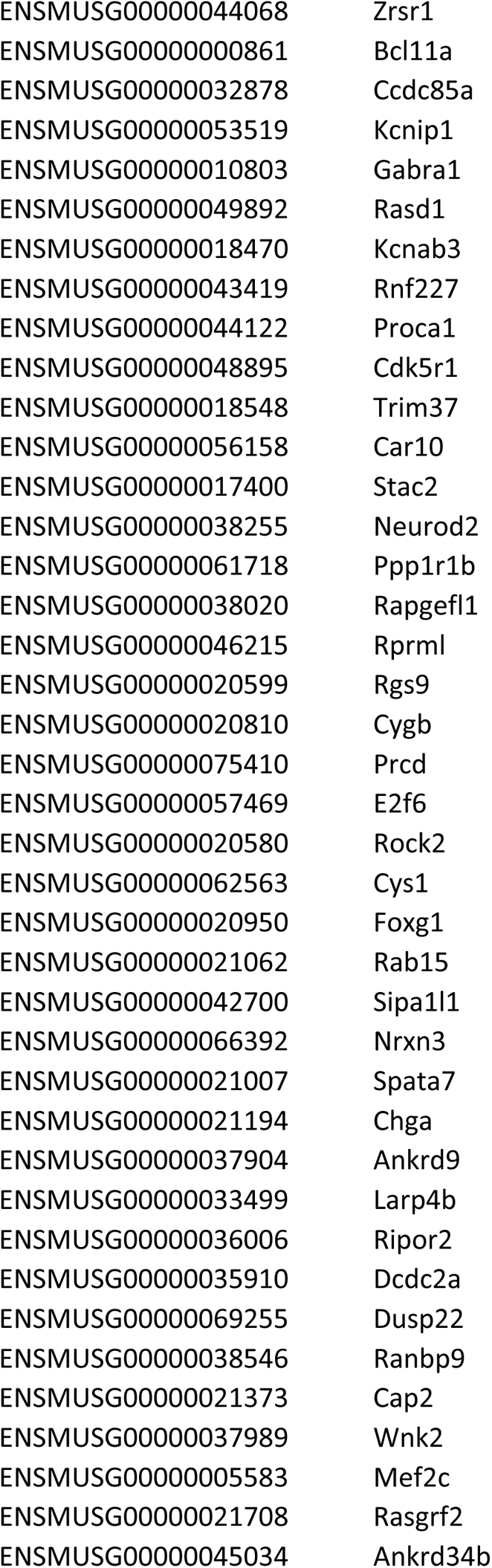
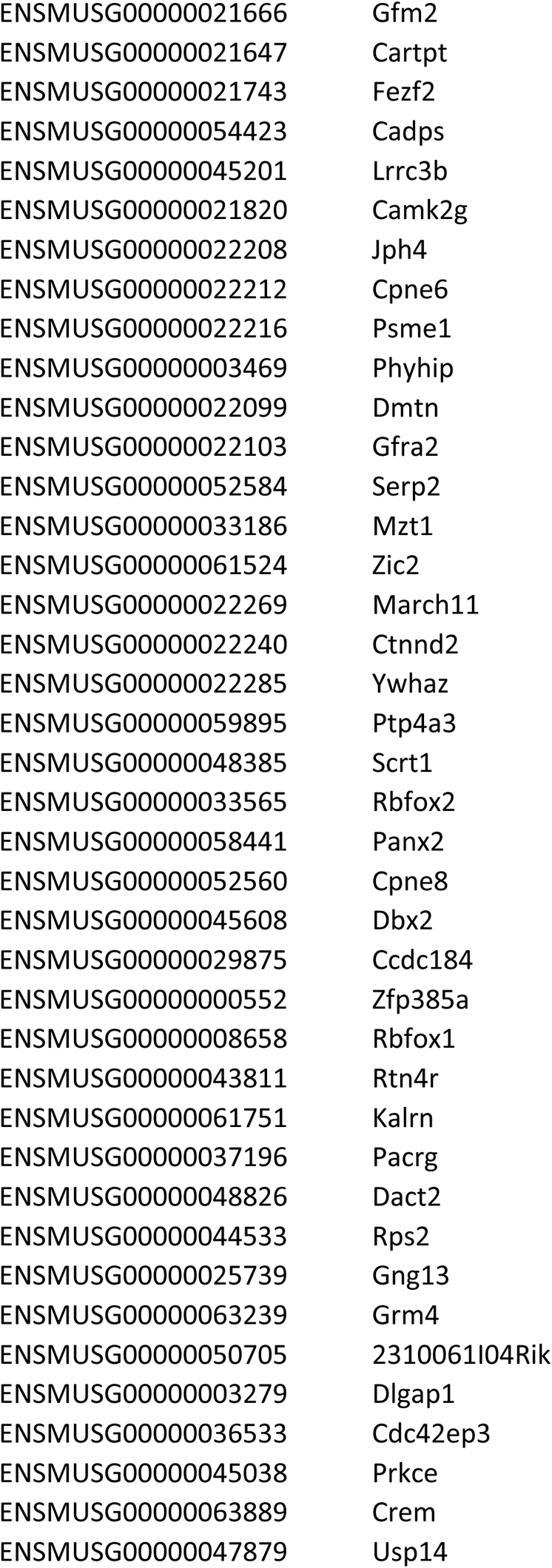
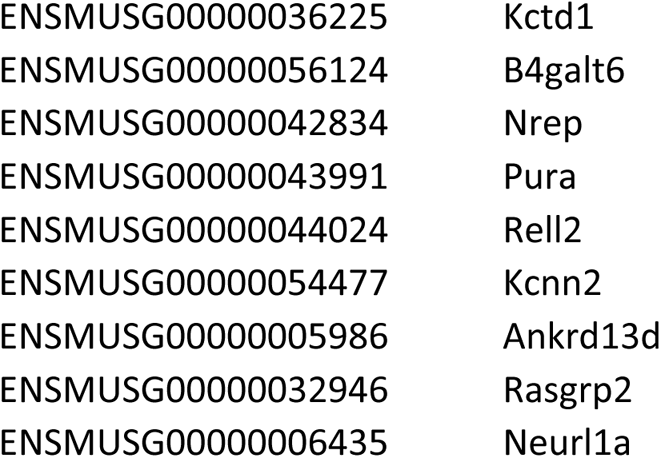
List of genes that are putative shared 4E-BP1 and eIF4E translational targets in myelin

**Figure 6.**
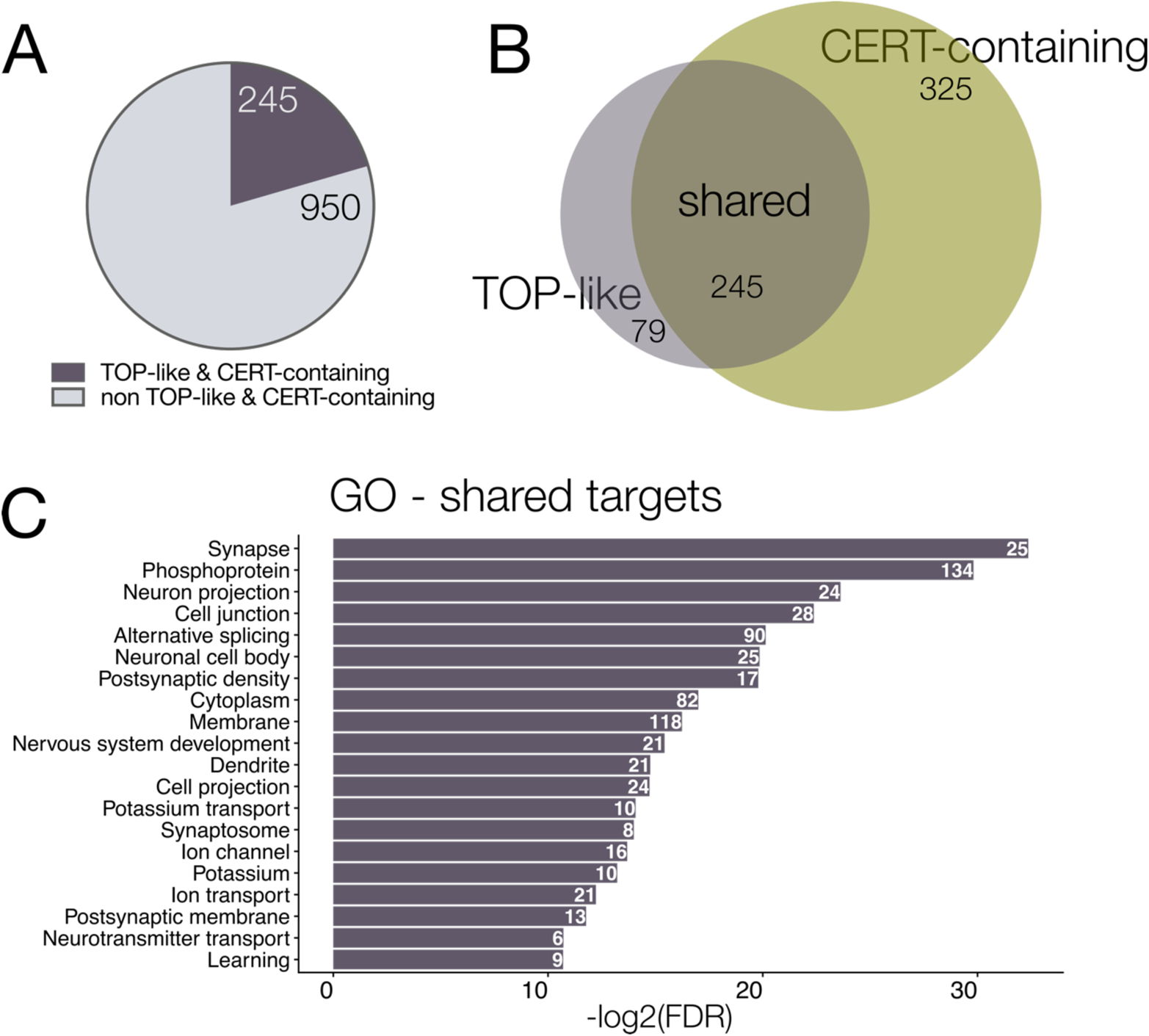
Identification of putative 4E-BP1 and eIF4E shared targets. (A) Percentage of myelin-resident mRNAs containing both a TOP-like motif and CERT sequence in their 5’UTRs. (B) Schematic representing the proportion of shared and unique translational targets between 4E-BP1 and eIF4E. (C) Gene ontology analysis of 4E-BP1 and eIF4E shared targets. Top 20 terms shown, ordered most to least significant by −log2 of false discovery rate. Counts correspond to number of genes associated with each term.

### Cap-dependent translation regulated by Akt-mTOR signaling drives myelin sheath growth

Our data show that a downstream translational regulator of Akt-mTOR signaling localized to myelin and hundreds of myelin-localized transcripts contain *cis*-regulatory motifs that confer their translation sensitive to Akt-mTOR signaling. Does Akt-mTOR-dependent protein translation drive myelination? To assess this, we performed cell-specific manipulation of the downstream translational regulators 4E-BP1 and eIF4E. To first test if 4E-BP1 function is required for myelin development, we overexpressed a human allele of 4E-BP1 where threonines 37 and 46, which are obligate mTOR phosphorylation sites, were mutated to alanine (4EBP1^T37/46A^). These mutations render the protein unable to be phosphorylated by mTORC1 and released from its translational repressor state (Gingras et al., 1999), thereby inhibiting 4E-BP1-mediated translation (Figure 7A). Using the Tol2-transgenesis system, we transiently expressed mNeonCAAX, a wild-type allele of human 4E-BP1 (4EBP1^WT^) and mNeonCAAX, or 4EBP1^T37/46A^ and mNeonCAAX in oligodendrocyte lineage cells of wild-type animals using *sox10* regulatory DNA. Overexpression of 4EBP1^WT^ did not change myelin sheath length, number, or cumulative length of myelin made when compared to wild-type controls (Figure 7D-F). However, when mTOR-4E-BP1 signaling was perturbed by overexpression of the 4EBP1^T37/46A^ allele, oligodendrocytes generated myelin sheaths that were ~35% shorter than those of wild-type larvae or those expressing 4EBP1^WT^ (Figure 7D). Additionally, cells expressing 4EBP1^T37/46A^ produced fewer myelin sheaths than those of controls (Figure 7E). Together, the reduction in myelin sheath length and number resulted in a 50% reduction in the total length of myelin produced by cells expressing 4EBP1^T37/46A^ (Figure 7F). From these data, we conclude myelin development requires 4EBP1-dependent translation.

**Figure 7.**
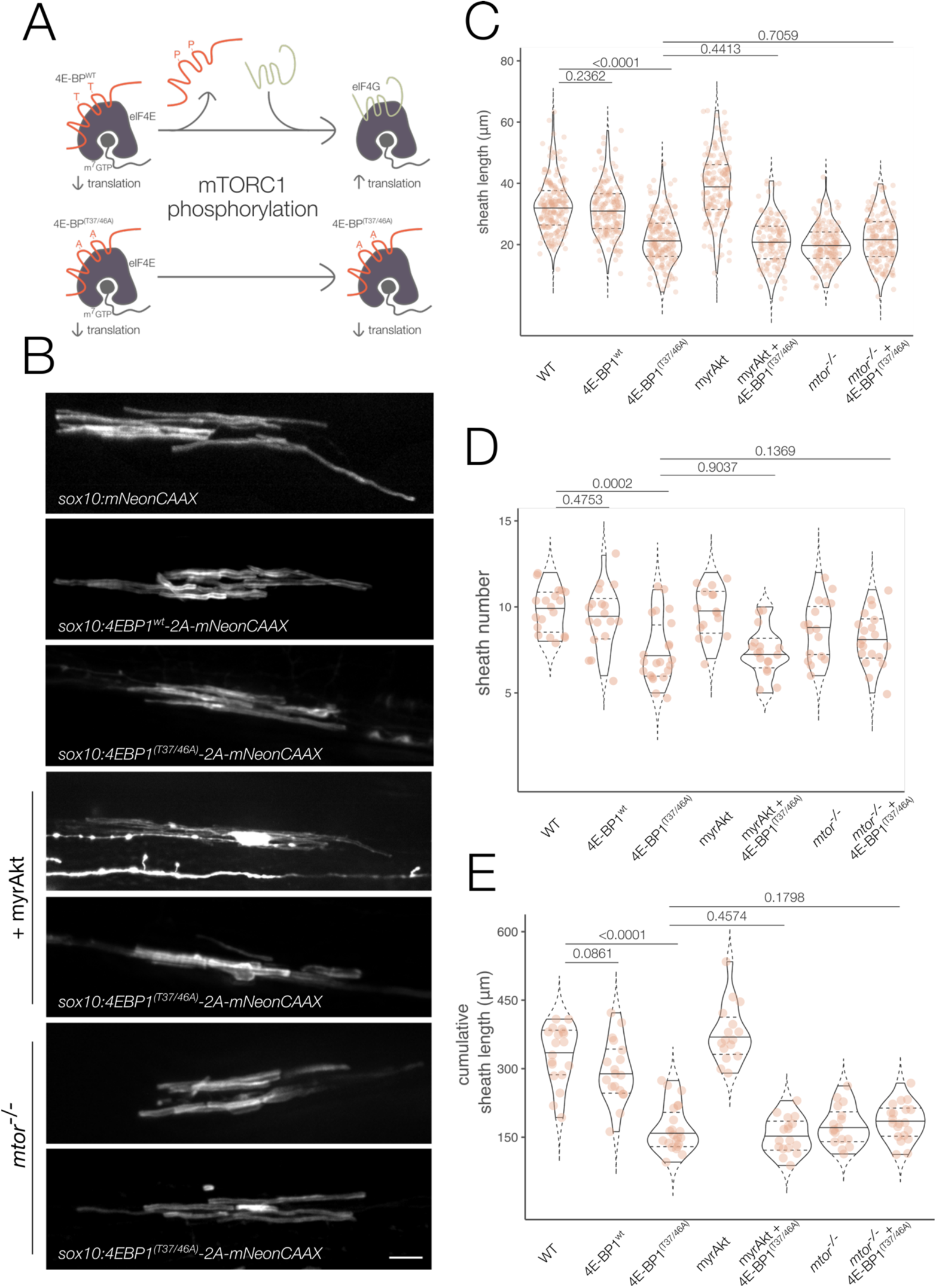
4E-BP1-dependent translation downstream of Akt-mTOR signaling promotes myelination. (A) Schematic illustrating the mechanism by which 4E-BPs promote translation initiation and the function disruption strategy. (B) Representative images of single 5 dpf spinal cord oligodendrocytes expressing controls or 4E-BP1 manipulation with or without simultaneous mTOR pathway manipulations. Scale bar, 10µm. Quantification of myelin sheath length (C), number (D), or cumulative myelin sheath length (E). Violin plot solid lines represent median and dashed lines represent 25^th^ and 75^th^ percentiles. Overall significance was assessed using the Kruskal-Wallis test: sheath length, p < 1×10^−12^, h = 458.4; sheath number, p = 8.62×10^−6^, h = 33.44; cumulative sheath length, p <1×10^12^, h = 86.76. Individual data points are represented by orange dots. Sample sizes: WT - 19 fish, 19 cells, 190 sheaths; 4EBP1^wt^ – 18 fish, 18 cells, 169 sheaths; 4EBP1^T37/46A^ – 22 fish, 22 cells, 179 sheaths; myrAkt – 16 fish, 16 cells, 155 sheaths; 4EBP1^T37/46A^ + myrAkt – 17 fish, 17 cells, 126 sheaths; *mtor^−/−^* - 17 fish, 17 cells, 148 sheaths; 4EBP1^T37/46A^ + *mtor^−/−^* - 20 fish, 20 cells, 154 sheaths. P values on figure are pairwise Mann-Whitney tests with Bonferroni-Holm correction for multiple comparisons.

As a direct test of whether the Akt-mTOR pathway requires 4E-BP1 function to drive myelination, we simultaneously manipulated Akt-mTOR signaling and 4E-BP1. To drive Akt-mTOR pathway activity in oligodendrocytes, we created the construct *sox10:mScarletCAAX-P2A-myrAkt.* Using the Tol2-transgenesis system, we expressed either *sox10:mScarletCAAX-P2A-myrAkt* alone or in combination with 4EBP1^T37/46A^ and mNeonCAAX in wild-type larvae. Simultaneous expression of myrAkt and 4EBP1^T37/46A^ resulted in a ~35% reduction in average myelin sheath length (Figure 7D). This was accompanied by a decrease in myelin sheath number (Figure 7E) and resulted in a nearly 50% reduction in the total length of myelin produced (Figure 7F). Individual and cumulative myelin sheath length as well as number produced by cells expressing both myrAkt and 4EBP1^T37/46A^ was not different than those of cells expressing 4EBP1^T37/46A^ alone (Figure 3D-F). To test if mTOR and 4EBP1 function in a common pathway to promote myelin development, we performed ensheathment analyses in *mtor* mutant larvae expressing either mNeonCAAX alone or 4EBP1^T37/46A^ and mNeonCAAX. Myelin sheath length, number, and cumulative length of myelin were unchanged in wild-type larvae expressing 4EBP1^T37/46A^ alone, *mtor* mutants, or *mtor* mutant larvae expressing 4EBP1^T37/46A^ (Figure 3D-F). From these data, we conclude that 4E-BP1-mediated translation downstream of Akt-mTOR signaling is required for proper myelin development.

Because our 4E-BP1^T37/46A^ data showed that dampening cap-dependent translation inhibits myelin development, we tested if increasing cap-dependent translation would drive myelin sheath elongation. Constitutive overexpression of eIF4E in the mouse brain exaggerates cap-dependent translation that results in alterations to synaptic morphology and physiology and animal behavior (Gkogkas et al., 2013; Santini et al., 2013). To test the role of eIF4E-mediated translation in our system, we expressed human eIF4E and mNeonCAAX in oligodendrocytes by Tol2-transgenesis. Oligodendrocytes expressing human eIF4E had, on average, a 10% increase in the average length of myelin sheaths (Figure 8B). This was accompanied by no change in the number of myelin sheaths produced (Figure 8C). Cumulatively, cells expressing human eIF4E produced a greater length of myelin sheaths (Figure 8D), but this did not reach statistical significance. Together, these data indicate that exaggerated protein translation alone is capable of driving myelin sheath elongation.

**Figure 8.**
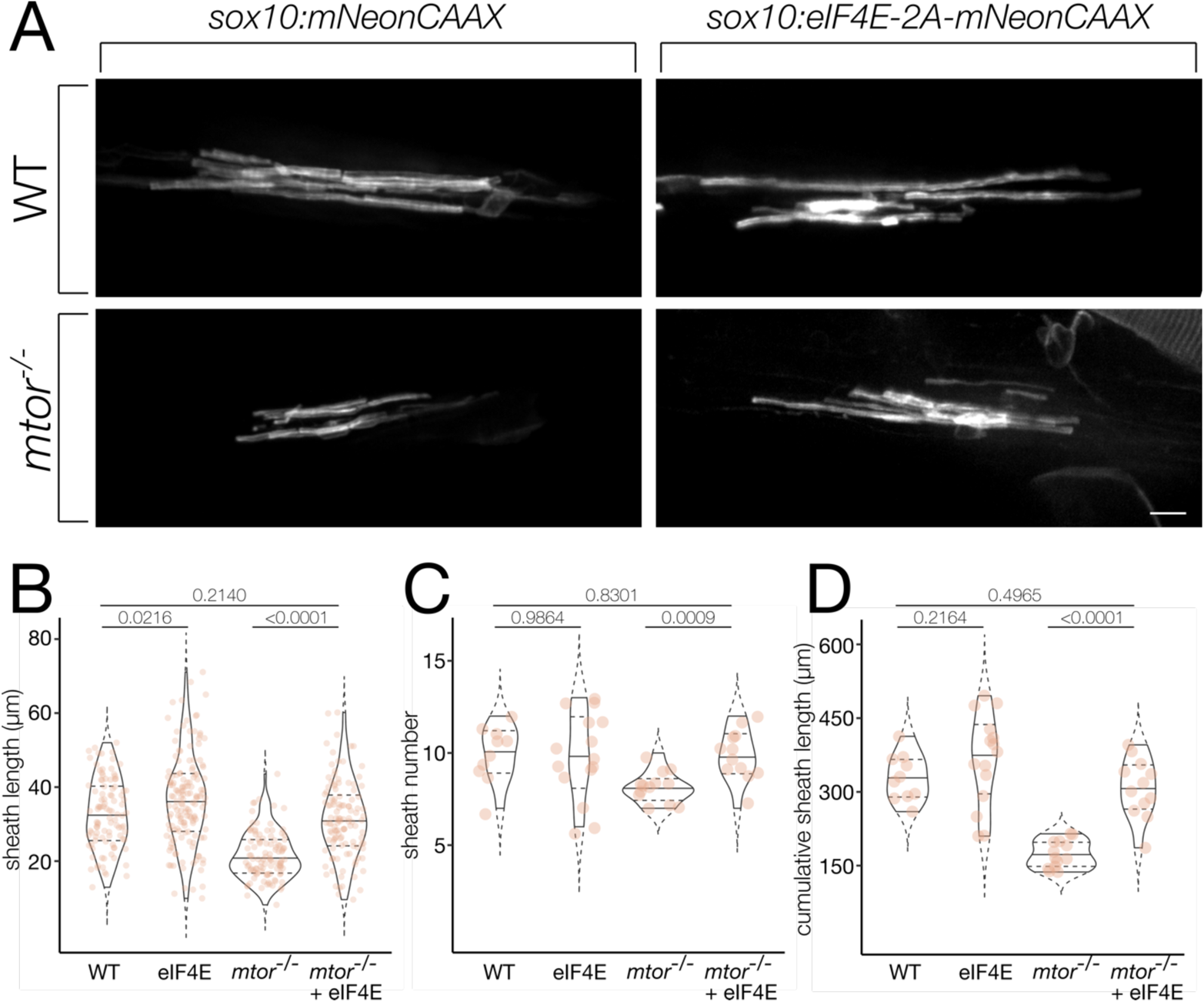
Upregulated eIF4E-mediated translation rescues mTOR loss-of-function. (A) Representative images of 5 dpf spinal cord oligodendrocytes in wild-type (WT) or *mtor* mutant larvae expressing human eIF4E or controls. Scale bar, 10µm. Quantification of myelin sheath length (B), number (C), and cumulative myelin sheath length (D). Violin plot solid lines represent median and dashed lines represent 25^th^ and 75^th^ percentiles. Overall significance was assessed by the Kruskal-Wallis test: sheath length, p < 1×10^−12^, h = 120.3; sheath number, p = 0.0069, h = 12.15; cumulative sheath length, p = 3×10^−6^, h = 28.44. Orange dots represent individual data points. Sample sizes: WT – 10 fish, 10 cells, 100 sheaths; eIF4E – 15 fish, 15 cells, 150 sheaths; *mtor^−/−^* - 13 fish, 13 cells, 104 sheaths; *mtor^−/−^* + eIF4E – 12 animals, 12 cells, 118 sheaths. P values on figure are pairwise Mann-Whitney tests with Bonferroni-Holm correction for multiple comparisons.

Our results showing 4E-BP1-dependent translation promotes myelination led us to conclude that translation initiation downstream of Akt-mTOR is required for myelin sheath elongation. As a direct test of this possibility, we exaggerated translation in oligodendrocytes of *mtor* mutant larvae by expressing human eIF4E and mNeonCAAX using Tol2 transgenesis and *sox10* regulatory elements. If translation downstream of mTOR signaling promotes myelination, we predicted *mtor* mutant myelin sheath length and number deficits would be rescued upon exaggerated translation achieved by eIF4E overexpression. Indeed, eIF4E overexpression in oligodendrocytes of *mtor* mutant larvae restored average myelin sheath length (Figure 8B), number (Figure 8C), and cumulative myelin sheath length (Figure 8D) to wild-type levels. These experiments provide strong evidence that protein translation regulated by mTOR signaling is required for oligodendrocytes to produce proper length and number of myelin sheaths.

## DISCUSSION

During development, oligodendrocytes ensheath axons with myelin to support nervous system function. The Akt-mTOR pathway is a key regulator of radial myelin sheath growth, but our understanding of how this pathway promotes myelin sheath elongation and the downstream signaling events it regulates to promote myelination remain limited.

By inhibiting 4E-BP1 function we found that protein translation initiated by Akt-mTOR signaling drives myelin sheath growth. Previous studies have shown that Akt-mTOR signaling in oligodendrocytes promotes both transcription and translation of some myelin genes and proteins (Flores et al., 2008; Narayanan et al., 2009; Tyler et al., 2009; Tyler et al., 2011; Bercury et al., 2014; Lebrun-Julien et al., 2014; Wahl et al., 2014; Zou et al., 2014; Kearns et al., 2015) and translation only of other myelin proteins (Tyler et al., 2009; Tyler et al., 2011; Bercury et al., 2014; Lebrun-Julien et al., 2014; Wahl et al., 2014). Consequently, the independent roles that transcription and translation downstream of Akt-mTOR signaling play in regulating myelin development are unclear, and investigations into how mTOR-dependent translation specifically promotes myelination have been limited. Ultrastructural analysis of myelin sheath thickness of 2-month-old 4E-BP1/2 knockout mice showed no change compared to wild-type siblings (Lebrun-Julien et al., 2014). From our data showing inhibited 4E-BP1-dependent translation results in shorter and fewer myelin sheaths, we would have predicted that increasing translation by removing 4E-BP1/2 function would lead to thicker myelin. However, it is possible that changes in myelin sheath thickness were apparent at younger ages not analyzed, consistent with the role of mTOR signaling in developmental myelination, or that mTOR signaling regulates different mechanisms to promote radial versus lateral growth of myelin sheaths.

Consistent with the role of protein translation driving myelination downstream of Akt-mTOR signaling, we found that eIF4E overexpression cell autonomously rescued myelin sheath length and number deficits observed in mTOR loss-of-function mutants. eIF4E represents a point of convergence between Akt-mTOR signaling and the MAPK/ERK pathway. Phosphorylation of eIF4E at Ser201 by Mnk1 directly regulates eIF4E cap-binding activity downstream of ERK signaling (Waskiewicz et al., 1997). Pharmacological treatment of mice with the mTOR inhibitor rapamycin led to an upregulation of phospho-ERK1/2 in oligodendrocytes of the corpus callosum (Dai et al., 2014), indicating mTOR signaling may negatively regulate ERK signaling during development. To date crosstalk between mTOR and ERK signaling during myelination have proposed Insulin Receptor Substrate-1 (Dai et al., 2014) or p70S6K (Michel et al., 2015) as points of convergence. Our data raise the possibility that eIF4E represents yet another integration point of these pathways, and that upregulation ERK signaling during development could overcome myelin sheath elongation deficits seen upon mTOR loss-of-function.

Do 4E-BP1 and eIF4E promote myelin sheath growth by targeting translation of the same transcripts? By mining the myelin transcriptome for cis-regulatory elements that confer translational control by 4E-BP1 or eIF4E, we identified populations of mRNAs whose translation may be regulated by either 4E-BP1 or eIF4E alone or the two in tandem. In support of our identification of different subsets of genes, yeast genetic studies show that 4E-BP family members can regulate both eIF4E-dependent and independent translation (Castelli et al., 2015). The myelin transcriptome is composed of hundreds of mRNAs with diverse functions (Thakurela et al., 2016). Gene ontology analysis revealed that the subsets of mRNAs we identified as potentially regulated by 4E-BP1 and/or eIF4E have distinct functions, raising the possibility that 4E-BP1 and eIF4E regulated transcripts have different functional consequences for myelin development. Another possibility is that 4E-BP1 and eIF4E function provide a level of regulation of selective translation based on extracellular signals. Additionally, interactions with RNA binding proteins such as FMRP, which localizes to and promotes myelination (Doll et al., 2020), could provide complex translational regulatory networks in myelin sheaths.

Our data provide preliminary support for a model wherein mTOR signaling in myelin sheaths promotes selective translation of myelin-resident mRNAs. Such a model has been tested in other cell types of the nervous system. In cultured cortical neurons, brain-derived neurotrophic factor (BDNF) stimulation promoted the phosphorylation of mTOR downstream targets 4E-BP and p70S6K in dendrites and drove neurite-localized translation of Arc and CaMKII in an mTOR-dependent manner (Takei et al., 2004). Our bioinformatic analyses revealed some CaMKII subunits as putative 4E-BP1 translational targets in myelin. Genetic studies in mice showed that both some CaMKII subunits (Waggener et al., 2013) and BDNF (Xiao et al., 2010; Lundgaard et al., 2013; Wong et al., 2013; Fletcher et al., 2018) promote central nervous system myelination, raising the possibility that BDNF coordinates new myelin synthesis by mTOR-dependent translation.

This model of mTOR signaling activating the translation of specific myelin-resident transcripts in a context-dependent manner is predicated on localized signaling and translation in myelin sheaths. In support of this possibility, we found a 4E-BP1 fusion protein and *mbpa* mRNA localized to myelin during development. Additionally, a reporter for PIP_3_ formation, the phospholipid that activates Akt, and phosphorylated Akt localized to the membrane of cultured oligodendrocytes (Snaidero et al., 2014). Moreover, in oligodendrocyte-dorsal root ganglion co-cultures, action potentials induced de novo translation of an *Mbp*-fluorescent reporter construct (Wake et al., 2011). In contrast to our proposed mechanism, these studies attributed *Mbp* translational regulation to Fyn kinase (Wake et al., 2011), however it is possible that differences in translational control could be dependent upon context or developmental age or even specific transcripts being regulated by different signaling mechanisms. Future work into this topic should aim to identify the spatial and temporal dynamics of signaling networks in nascent myelin sheaths as well as validate the local translation of myelin-resident mRNAs in vivo.

## Acknowledgements

We thank the Appel lab for helpful discussions, Drs. Caleb Doll and Alexandria Hughes for constructive comments on the manuscript, Dr. Katie Yergert for experimental guidance, and Rebecca O’Rourke for help with bioinformatics analysis. This work was supported by US National Institutes of Health (NIH) grant R01 NS095670 and a gift from the Gates Frontiers Fund to B.A. K.N.F-S. was supported by NIH F31 NS115261 and NIH T32 NS099042. The University of Colorado Anschutz Medical Campus Zebrafish Core Facility was supported by NIH grant P30 NS048154.

